# PASER for automated analysis of neural signals recorded in pulsating magnetic fields

**DOI:** 10.1101/739409

**Authors:** Terence Brouns, Tansu Celikel

## Abstract

Thanks to the advancements in multichannel intracranial neural recordings, magnetic neuroimaging and magnetic neurostimulation techniques (including magnetogenetics), it is now possible to perform large-scale high-throughput neural recordings while imaging or controlling neural activity in a magnetic field. Analysis of neural recordings performed in a switching magnetic field, however, is not a trivial task as gradient and pulse artefacts interfere with the unit isolation. Here we introduce a toolbox called PASER, Processing and Analysis Schemes for Extracellular Recordings, that performs automated denoising, artefact removal, quality control of electrical recordings, unit classification and visualization. PASER is written in MATLAB and modular by design. The current version integrates with third party applications to provide additional functionality, including data import, spike sorting and the analysis of local field potentials. After the description of the toolbox, we evaluate 9 different spike sorting algorithms based on computational cost, unit yield, unit quality and clustering reliability across varying conditions including self-blurring and noise-reversal. Implementation of the best performing spike sorting algorithm (KiloSort) in the default version of the PASER provides the end user with a fully automated pipeline for quantitative analysis of broadband extracellular signals. PASER can be integrated with any established pipeline that sample neural activity with intracranial electrodes. Unlike the existing algorithmic solutions, PASER provides an end-to-end solution for neural recordings made in switching magnetic fields independent from the number of electrodes and the duration of recordings, thus enables high-throughput analysis of neural activity in a wide range of electro-magnetic recording conditions.

## 1 Introduction

Magnetic control of neural activity is an emerging technology that promises non-invasive brain stimulation [1, 2]. Because magnetic fields can penetrate the body largely undisturbed, magnetic stimulation can be performed without implanting a probe in the brain [3, 4]. Magnetic neural stimulation can be combined with optical and electrical methods for multimodal observation and control of neural circuits. It offers faster temporal resolution than other noninvasive brain stimulation techniques (e.g. ultrasound, transcranial magnetic stimulation) [1, 5].

Electrical recording in a switching magnetic field is nonetheless not a trivial task due to electromagnetic interference and the associated electrical, including gradient and pulse, artefacts [6, 7]. Gradient artefacts are bipolar signals that are several orders of magnitude larger than the neural signals, caused by a switch in magnetic field. Because they are transients, triggered externally, they could be largely removed by subtracting an average gradient artefact trace from broadband neural signals [8]. Alternatively they could be isolated using various blind and supervised source separation techniques [9–12], all of which have been previously applied to human electroencephalographic data [13–15]. Pulse artefacts are smaller than gradient artefacts and vary primarily based on blood motion [7]. It has been suggested that pulse artefacts are caused by a) electromagnetic induction [16], b) the Hall effect which is due to the movement of conductive fluid in a static magnetic field, inducing electrical potentials [17], and c) the pulsatile motion of the living tissue, caused (at least in part) by the blood flow dynamics in vasculature [7]. Although various independent component analysis (ICA) methods [6,14,18,19] have been developed to remove pulse artefacts, ICA decomposition requires source to be stationary, hence it performs poorly when subjects are in motion in respect to the conductor (electrode). Moreover, in experimental conditions where the distance between a conductor and electromagnet varies across trials, gradient artefacts would vary significantly, requiring detection and removal of each artefact separately. For these reasons chronic neural recordings in a pulsing magnetic field requires novel signal processing approaches. Here we introduce a toolbox called PASER, Processing and Analysis Schemes for Extracellular Recordings, that performs automated denoising and quality control of electrical recordings. PASER is written in MATLAB and modular by design. The current version integrates third party applications to provide additional functionality, including with OpenEphys [20] for data import, KiloSort [21] for spike sorting and FieldTrip [22] for the analysis of local field potentials. PASER also provides quantitative measures of unit quality based on statistical criteria for objective classification of spike sorting results. The toolbox is distributed freely via github.com/DepartmentofNeurophysiology/Paser and should facilitate high-throughput, automated analysis of invasive neural recordings made under challenging conditions, including during electromagnetic interference.

## 2 Methods and Results

Figure 1 illustrates the off-line data processing pipeline implemented by PASER. Broadband extracellular signals are measured typically by a polytrode and is sampled at ∼30 kHz [23–28]. We show a tetrode in the diagram for illustrative purposes, but higher-channel probes are also supported by the toolbox. What follows are descriptions of the various processing steps carried out by PASER and that are presented in Figure 1. Note that many of these stages likely to have alternative solutions, which is why PASER has a modular pipeline, so that each stage can easily be modified as methodology develops further. The processing steps are under user control through global parameters. In the text, we report default parameter values, but the user can provide their own combination of parameter values in an external script that can be loaded by the toolbox as the pipeline can be run in a fully automated manner.

**Figure 1:**
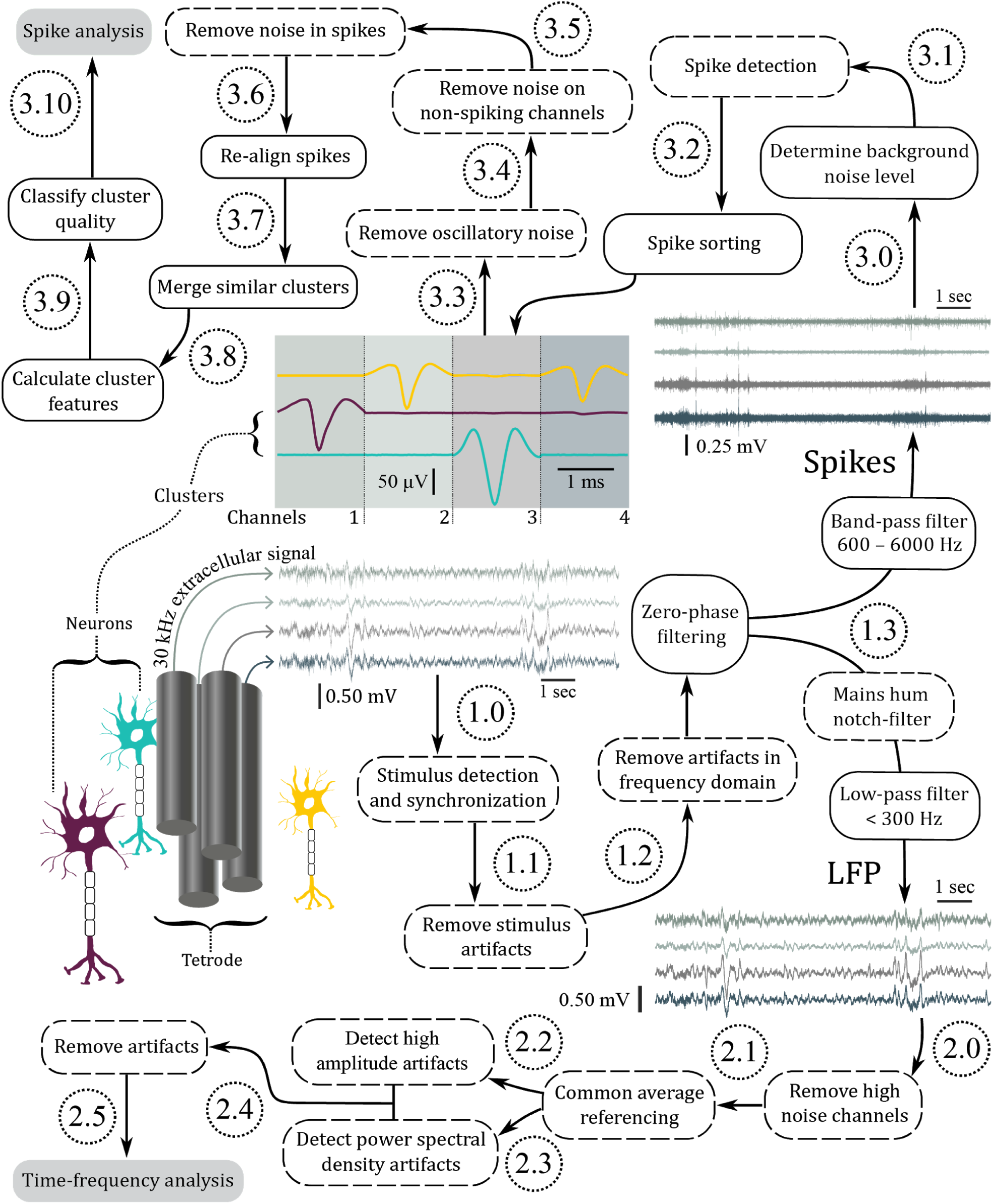
The flow-chart of PASER. The pipeline takes the raw broadband extracellular signals measured by any number of electrodes or polytrodes (here shown as a tetrode), removes electromagnetic and other stimulus related artefacts before isolating units and LFP data for analysis. In the figure, steps are represented by boxes, where a solid outline indicates a mandatory step while a dashed outline is optional, but often recommended. See main text for descriptions of each step.

After the description of the processing steps, we evaluate 9 different spike sorting algorithms based on computational cost, unit yield, unit quality, and clustering reliability (robustness) when signals are disturbed through self-blurring and noise-reversal [29] and implement one of the best performing spike clustering methods, i.e. KiloSort [21], in PASER to provide the end-user a fully automated pipeline.

### 2.1 Pre-processing

In the first section of the pipeline, we carry out operations on the raw signal to eliminate noise and extract metadata, before splitting the pipeline into the spike and LFP processing branches. To respect the memory capacity of typical workstations, data is partitioned and processed individually.

#### Step 1.0: Stimulus detection and synchronization

Stimulus delivery, let it be sensory or through brain stimulation, is a popular choice in experimental designs that aim to address the causality between neural activity and (network and organismal) behavior. In these experiments the timing of each stimulus is synchronized with the clock of the extracellular measurement in order to capture the neurophysiological response. Since this synchronization procedure is experiment dependent, PASER allows the user to provide stimulus information. The detected stimulus timestamps are subsequently stored as metadata in the final output file for use in the analysis.

#### Step 1.1: Stimulus-induced artifact removal

During electrical recordings performed in pulsating magnetic fields, changes in the gradient cause a (large) deflection in the voltage time-series due to electromagnetic induction (see Figure 2). If these transients are not removed, the data around them are dramatically distorted due to the appearance of prominent high-frequency ringing artifacts after band-pass filtering (see Step 1.3), ruling out any possibility of spike detection around the stimulus. To correct for this contamination, we remove the transients after detecting them using a derivative threshold of 10 times the median absolute deviation (MAD) and then translating the intermediate data segment bounded by the two transients so that it interpolates between start and end points, which conserves the data segment for further processing. Only spikes can be retrieved within the filtered signal, because significant low frequency contamination remains, so the LFP is lost due to linear interpolation.

**Figure 2:**
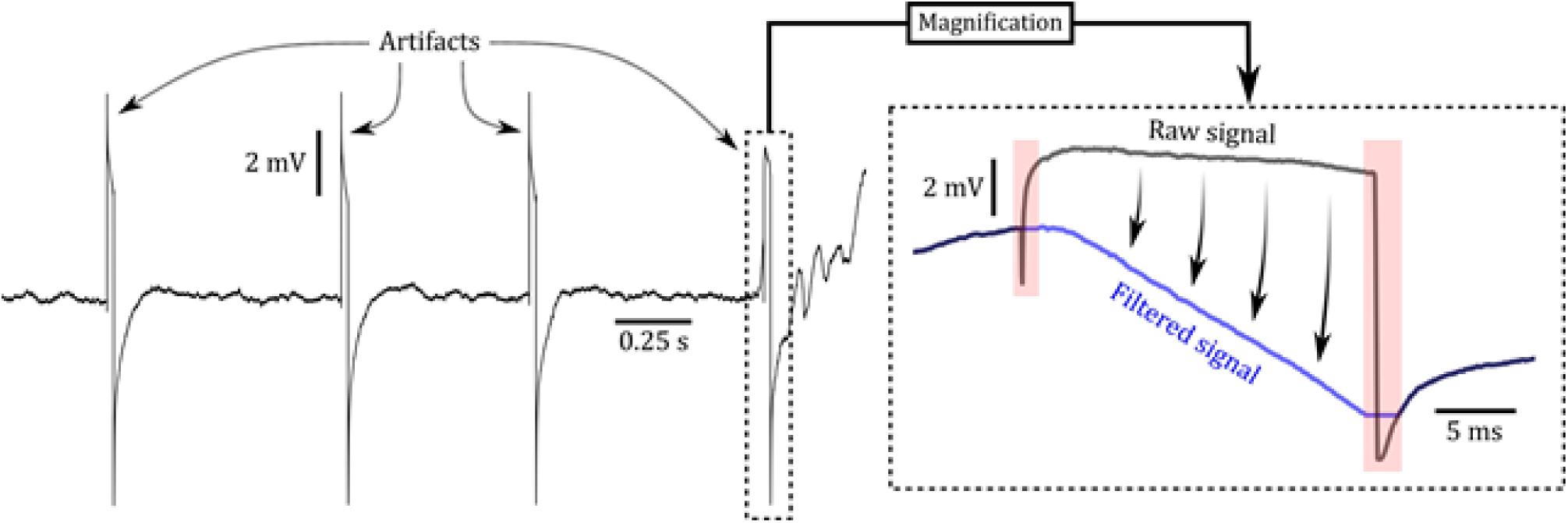
Magnetic stimulus artifact removal. The magnetic pulses produce large transient events in voltage time series due to electromagnetic induction. These peaks, if left intact, will introduce ringing artifacts upon band-pass filtering. The artifacts are thus removed by silencing the transients and then transforming the segment in-between. The transformation is denoted by the four arrows in the right plot. In this case, the resulting signal ends up having a large downward slope, but this low-frequency component is removed by band-pass filtering before spike detection.

#### Step 1.2: Frequency domain artifact removal

PASER can adaptively remove noise from external sources such as power lines and electronic oscillators by turning to the frequency domain. The neural signal is expected to have a power that is inversely proportional to the frequency (1/f), while external sources of noise will often exhibit narrow peaks in the power spectrum [30]. Fast Fourier transform (FFT) is applied to the raw data and outliers in the resulting spectrum are detected using a 5 MAD threshold, which are subsequently removed by setting them to zero and then applying inverse FFT to retrieve the filtered data.

#### Step 1.3: Zero-phase filtering

To separate the spikes from the LFP, we band-pass filter the signal between 600–6000 Hz to produce the data for spike detection, while a 300 Hz low-pass filter is used to form the LFP signal after filtering out the mains hum using a notch filter (50 Hz). Zero-phase filtering is applied, because phase nonlinearities have been shown to distort spike shapes and generate artificial spikes [31, 32].

### 2.2 Low-frequency signal processing

In this branch of the pipeline, we take the low-pass filtered local field potential signal and prepare it for time-frequency analysis by removing noise and artifacts.

#### Step 2.0: Remove high noise channels

Before further LFP processing, we remove channels that have a mean absolute amplitude that is three standard deviations greater than the overall mean of the other channels. This is done to avoid detecting an excessive amount of artifacts in the upcoming stages. Such channels often do not contain usable signals due to broken wiring in the electrodes or inadequate electrode impedance [33].

#### Step 2.1: Common average referencing

External noise, such as mechanical displacements and electromagnetic interference, will often occur simultaneously on neighbouring channels. We can reduce correlated noise by subtracting the average across all channels from each channel, a technique referred to as common average referencing (CAR) [30, 34]. Care should be taken when implementing CAR for probes with a relative low channel count, such as stereotrodes or tetrodes [35], given that the LFP signal could be strongly correlated across the channels of the probe, causing the CAR to remove relevant LFP signatures as well.

#### Step 2.2: Detect amplitude artifacts

Artifacts in extracellular recordings are frequently characterised as having significantly larger magnitudes and sharper signal transitions compared to the neural data of interest. For this reason, we classify data sections as artifacts if they have an absolute signal magnitude and/or derivative greater than an upper threshold of 6 MAD. These artifact regions are then demarcated using a lower threshold of 2 MAD (see Figure 3). The derivative is calculated using a step-size of 6 ms, which is a temporal resolution that was empirically found to correspond with the time-scale of the typical artifact ramp-up. Furthermore, we use the average signal across the channels of each probe to detect the artifacts on a probe-by-probe basis, since high-amplitude artifacts are generally shared across the channels of the same probe, but not across different probes.

**Figure 3:**
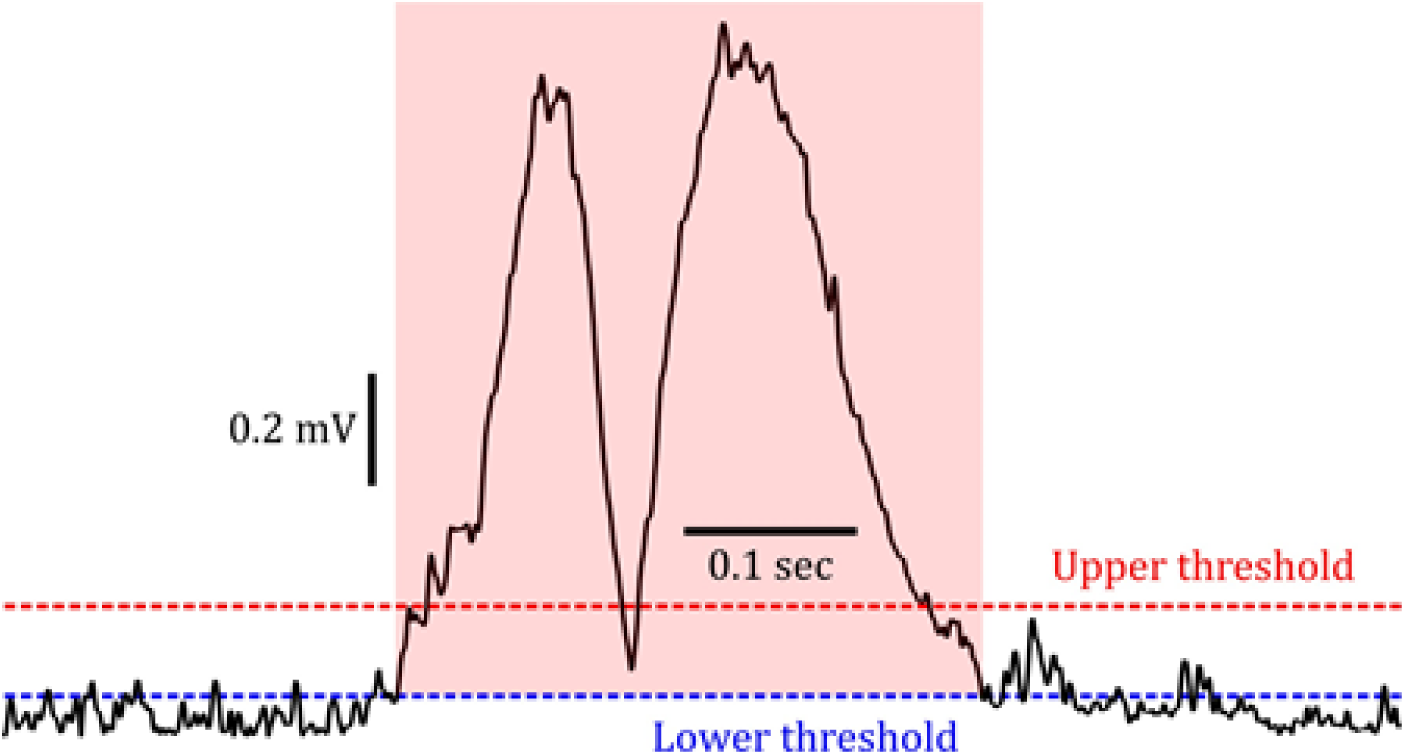
Method for elimination of high-amplitude artifacts in the LFP signal. Shown is the magnitude of an extracellular voltage trace over a short recording period, containing a brief artifact region. The artifacts are detected by the upper threshold (6 MAD), but are removed up to the point where they dip below the lower threshold (2 MAD). The data section that is deleted is indicated by the red shaded area. We use the absolute value of the signal, so only one pair of thresholds needs to be set. We classify a segment as an artifact if its PSD magnitude exceeds 3 MAD, where the MAD is calculated across all segments of the recording.

#### Step 2.3: Detect power spectral density artifacts

Artifacts can also be identified by their power spectral density (PSD). For example, a mechanical motion artifact will have high power signal bands that are spread across the whole frequency spectrum [36]. Therefore, we split the recording session into segments of 0.5 seconds with 0.25 seconds of overlap and calculate the PSD in every segment using Welch’s method for 64 frequencies evenly distributed between 10 and 100 Hz. We then normalize the PSD for each frequency by dividing by the average PSD for that frequency across all data segments and subsequently sum the PSDs across all frequencies. This results in a scalar value representing the PSD magnitude for each segment.

#### Step 2.4: Artifact removal

The artifact intervals detected in Step 2.2 & 2.3 are set to Not-a-Number (NaN) in the data, so these sections can be ignored during the analysis. Certain analyses, however, cannot be performed on data containing NaNs, so in those cases NaNs are substituted by zeros or an interpolation between neighbouring real numbers. Before NaN substitution, adjacent artifact intervals are joined together when there is less than 0.5 seconds between them, where we make the assumption that the intermediate region is probably contaminated by noise.

#### Step 2.5: Time-frequency analysis

To perform time-frequency analysis on the LFP data, the toolbox provides wrapper functions to call routines in the external MATLAB software toolbox FieldTrip [22]. These include the calculation of time-frequency representations using sliding time windows or the plotting of spectrograms and power spectra. Users can also create their own custom analysis scripts that take the processed LFP data as input and perform more sophisticated and detailed analyses for their particular experiment.

### 2.3 High-frequency signal processing: Action potential detection

In this branch of the pipeline, we take the band-pass filtered signal, detect the spikes contained within and associate them with their source neurons, which is followed by noise removal routines and quality control.

#### Step 3.0: Determine background noise level

For a number of stages during spike processing, it is necessary to have a reliable estimate of the background noise level, σ, for every channel. However, common measures such as the standard deviation or the root-mean-square of the signal are heavily influenced by high amplitude artifacts as well as the spike waveforms themselves, leading to an excessive magnitude of σ. A better estimate can be obtained by either calculating the MAD or the signal envelope, which are less sensitive to peaks in the data. The toolbox uses the signal envelope by default, which is given by the magnitude of the analytical signal calculated through Hilbert transform [37].

#### Step 3.1: Spike detection

By default PASER extracts a window of 1.5 ms in length around any negative peak in the signal that exceeds 3σ in any channel. In case of overlap between spike windows, we keep the spike that has the highest relative amplitude compared to σ. The resulting spike waveforms and corresponding timestamps are then stored for further processing.

#### Step 3.2: Spike sorting

Usually an electrode will measure the activity of multiple surrounding neurons, each producing a spike with a characteristic waveform. To assign each waveform to a “unit”, spike sorting is performed. Details of the spike sorting as well as a detailed comparison across 9 different toolboxes are provided below. For now, we simply note that spike sorting will output a number of distinct spike waveform clusters, where each cluster of spikes putatively belongs to a particular neuron. We will use the term unit rather than neuron, in accordance with the spike sorting literature, due to the fact that the clusters are the result of statistical inference [38].

#### Step 3.3: Remove oscillatory noise

One common observation in the spike sorting output is the presence of high-frequency oscillatory noise [39] that has been erroneously labelled as a spike. One example of this is shown in Figure 4. These waveforms are typified by repeating sinusoidal waves, unlike waveforms of physiological spikes, which allows us to use FFT to detect them. For each cluster of spikes and every channel individually, we calculate the FFT of its median waveform in time windows of approximately 7 ms and then find the maximum amplitude in the resulting power spectrum. If the largest peak exceeds 0.15 of the total spectral power, the channel is classified as a “ripple” and is masked with Gaussian white noise with standard deviation equal to σ. We typically calculate the median waveform, rather than the mean waveform, because the median is less susceptible to high-amplitude outliers (see Step 3.5).

**Figure 4:**
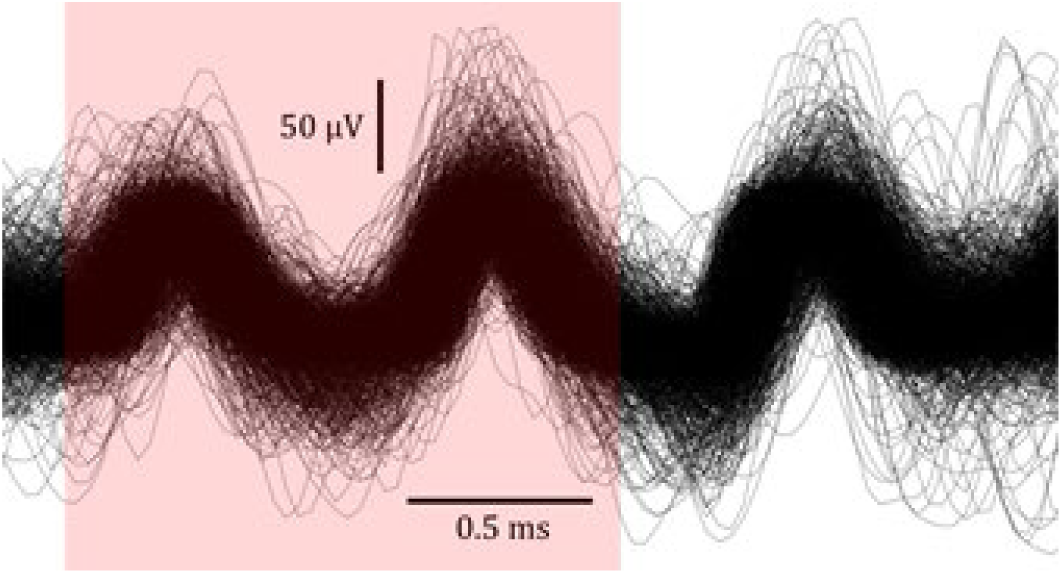
Example of high-frequency oscillatory noise. The red shaded region indicates the erroneously extracted spike waveform.

#### Step 3.4: Remove noise in non-spiking channels

When recording with polytrodes not every channel necessarily records action potentials. When we detect a spike on one or more channels, it would be preferable if the other channels are noise-free. This is particularly crucial when we have clusters that are mistakenly split up due to noise in non-spiking channels, but should in fact be merged together as these clusters collectively contain spikes that were fired by the same neuron. We can differentiate between noise and genuine spikes by comparing the median waveform in each channel with the median waveform in the largest amplitude channel, relative to σ.

Spikes of individual cells are often recorded simultaneously by several channels of the probe, where the largest amplitude waveform corresponds with the electrode that is closest to the neuron and smallest amplitude for the farthest [40]. Therefore, channels containing an extracellular spike waveform should be similar in shape, but not necessarily in magnitude. For this reason, we normalize the waveform in each channel by dividing the waveform by the median amplitude, so the magnitude in each channel should be approximately equal, centred around 1.0. We then calculate the mean-squared error (MSE) between each channel and the largest amplitude channel. If MSE > 0.05, the channel is masked by Gaussian white noise, similarly as in Step 3.3. Furthermore, the peak of the spike waveform should be approximately temporally aligned across the different channels in which it is observed, so we also calculate the difference (ΔT) in peak location between the largest amplitude channel and the other channels (see Figure 5). If ΔT > 0.15 ms, the channel is replaced by Gaussian white noise as well.

**Figure 5.**
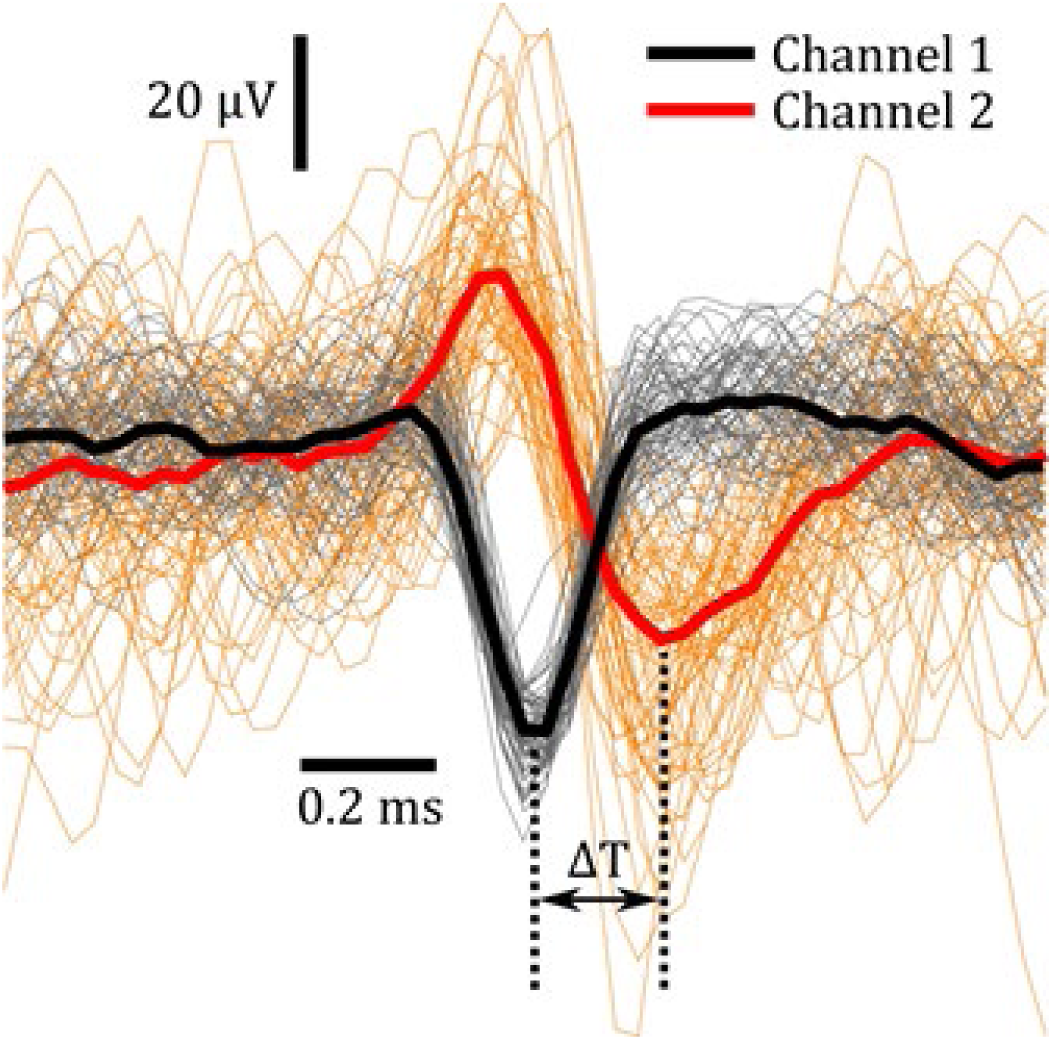
Different channels on the same probe are expected to show the same waveform shape when detecting a spike, but with varying amplitudes. Significant discrepancy between channels is indicative of noise contamination or multiple spikes overlapping. We detect this divergence by calculating the MSE between the waveforms and also the timing difference between the peaks (ΔT).

#### Step 3.5: Remove noise spikes

Spike sorting algorithms regularly insert artifacts that have action potential like waveform (see Figure 6). Due to their high amplitude, the artifacts are detected along with the physiological spikes, but are not put in their own separate cluster, because of their uniquely idiosyncratic waveforms and relatively low prevalence, so they are arbitrarily distributed over all clusters. These artifacts are dealt with by calculating the MSE between the cluster’s median waveform and every spike in that cluster, and then removing every spike with an MSE > 12σ. We also discard any spike that is detected at the location of the raw signal artifacts from Step 1.1.

**Figure 6.**
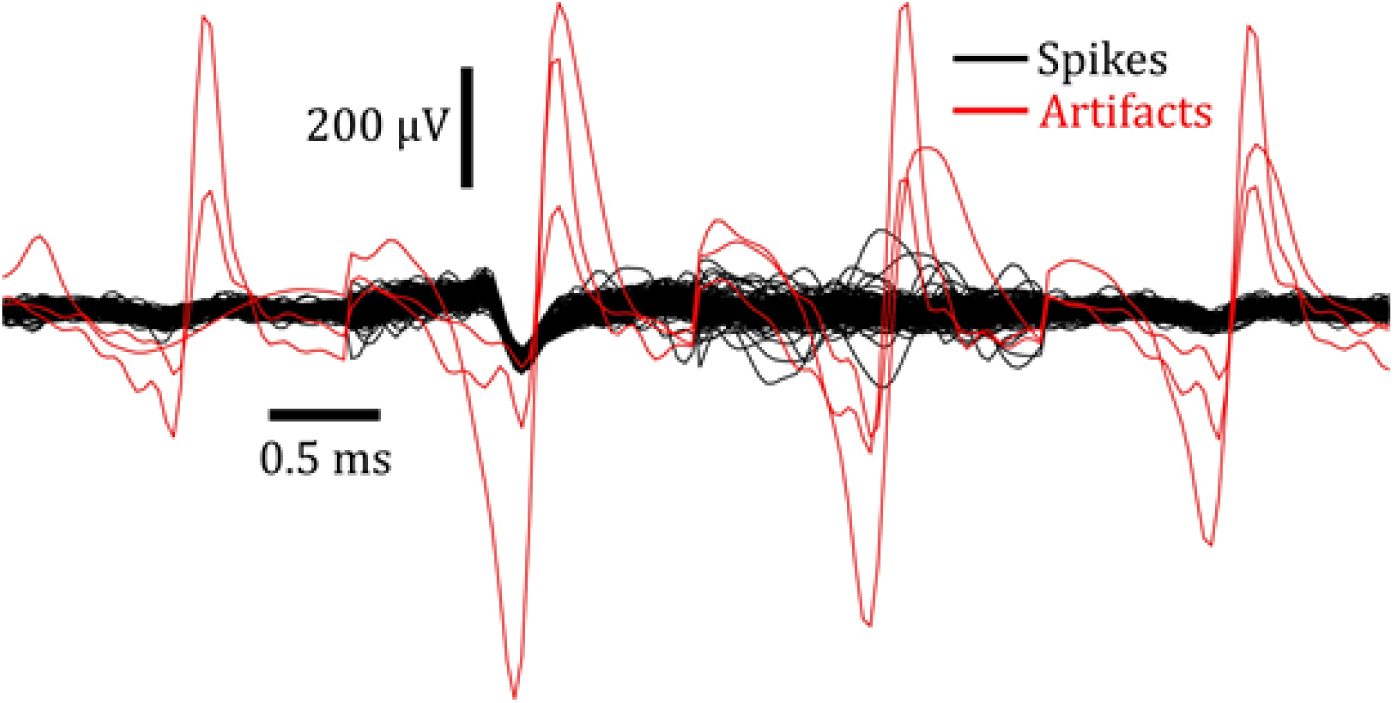
Spike waveforms of a single cluster overlaid on top of one-another, which are contaminated by high-amplitude artifacts (shown in red).

#### Step 3.6: Re-align spikes

After deleting noisy spikes, we (re-)align all remaining spikes to the position of the maximum amplitude (normalized by σ) in the cluster’s median waveform. This is crucial when merging spike clusters (see Step 3.7), because a slight temporal misalignment between the average waveforms of two clusters can prevent merging from happening, even though the two waveforms are otherwise identical and are highly likely to belong to the same unit.

#### Step 3.7: Merge similar clusters

To avoid under-estimation of action potential statistics, it is important that clusters corresponding to the same unit are merged. Merging clusters is typically a necessary post-hoc step, because spike sorting algorithms frequently produce a fixed number of clusters as their output and if this number is too large, then this leads to so-called over-clustering, i.e. two or more clusters corresponding to the same unit [33], which requires merging. Instead of merging, we could in theory prevent over-clustering from happening in the first place by tuning the parameters of the spike sorting algorithm more carefully, but over-clustering is actually desirable, because it helps with noise cancellation in the steps directly after spike sorting (see Steps 3.3–3.5). Furthermore, the opposite effect referred to as under-clustering, i.e. one cluster corresponding to multiple neurons, is harder to correct for, so we like to err on the side of over-clustering. In many spike sorting toolboxes, merging is done manually. However, we want this step to be automatic so it is more user-friendly and less subjective. We use the method employed by Yger et al. (2016) as our merging criterion, which is the normalized distance in principal component space between two clusters projected on the axis joining their two centroids [41].

#### Step 3.8: Calculate cluster features

Ideally, each resulting cluster corresponds to a single unique neuron. We refer to such clusters as isolated single units. Unfortunately, we often obtain clusters that only partially account for all the spikes of a neuron, or clusters that contain spikes of multiple neurons. We require objective quality control metrics that can be used to differentiate between these different types of units. Table 1 lists the most important cluster features that the toolbox calculates, which are subsequently used to score the quality of the unit (see Step 3.9). For descriptions of more cluster features, see the README file of the PASER toolbox at https://github.com/DepartmentofNeurophysiology/Paser/blob/master/README.md.

**Table 1:** List of cluster features that are computed by the toolbox and are utilized to quantify the unit quality. Each feature has a particular threshold, which a high-quality unit is expected to pass.

| Cluster feature name | Abbrev. | Threshold | Description |
| --- | --- | --- | --- |
| Spike count | $N_{spk}$ | $> 50$ | Total number of spikes in the cluster |
| Relative amplitude | $V_{rel}$ | $> 4\sigma$ | Median waveform amplitude divided by the spike detection threshold across all spikes in cluster |
| Absolute amplitude | $V_{abs}$ | $< 300 \mu V$ | Median waveform amplitude across all spikes in cluster |
| Peak-to-peak amplitude | $V_{p2p}$ | $< 400 \mu V$ | Voltage difference between maximum and minimum of median waveform |
| Refractory period violations | RPVs | $< 0.05$ | Fraction of spikes that occur within 1.5 ms after the previous spike in the cluster. |
| Subthreshold spikes | Subs | $< 0.10$ | Fraction of spikes in cluster that have an amplitude less than $3\sigma$. |
| Isolation quality | F1 score | $> 0.90$ | Harmonic mean of precision and recall of spikes in cluster determined by drifting t-distributions. |
| Temporal stability | $FS_{mse}$ | $< 4.00$ | Mean-squared error between empirical and theoretical Gaussian distribution of spike counts in overlapping intervals of 10 seconds. |
| Channel desynchronization | $\Delta T_{desync}$ | $< 0.25 ms$ | Peak location of cross-correlation between probe channels that are maximally dissimilar. |

#### Step 3.9: Classify cluster quality

The flow-chart of Figure 7 is used to provide clusters with different quality labels, depending on which and how many feature thresholds the cluster passes. Noise clusters are always excluded from the final output as these cannot be confidently viewed as neuronal activity, but rather fuzzy background activity at best. During the analysis, the user can decide which of the other three classes of units should be included in the analysis, which is dependent on the experimental question under investigation. The user is also free to ignore these quality classifications and instead rely on a custom classification routine based on different combinations of cluster features supplied by the toolbox as detailed in Table 1. For an in-depth quality control discussion see Section 3.2.

**Figure 7:**
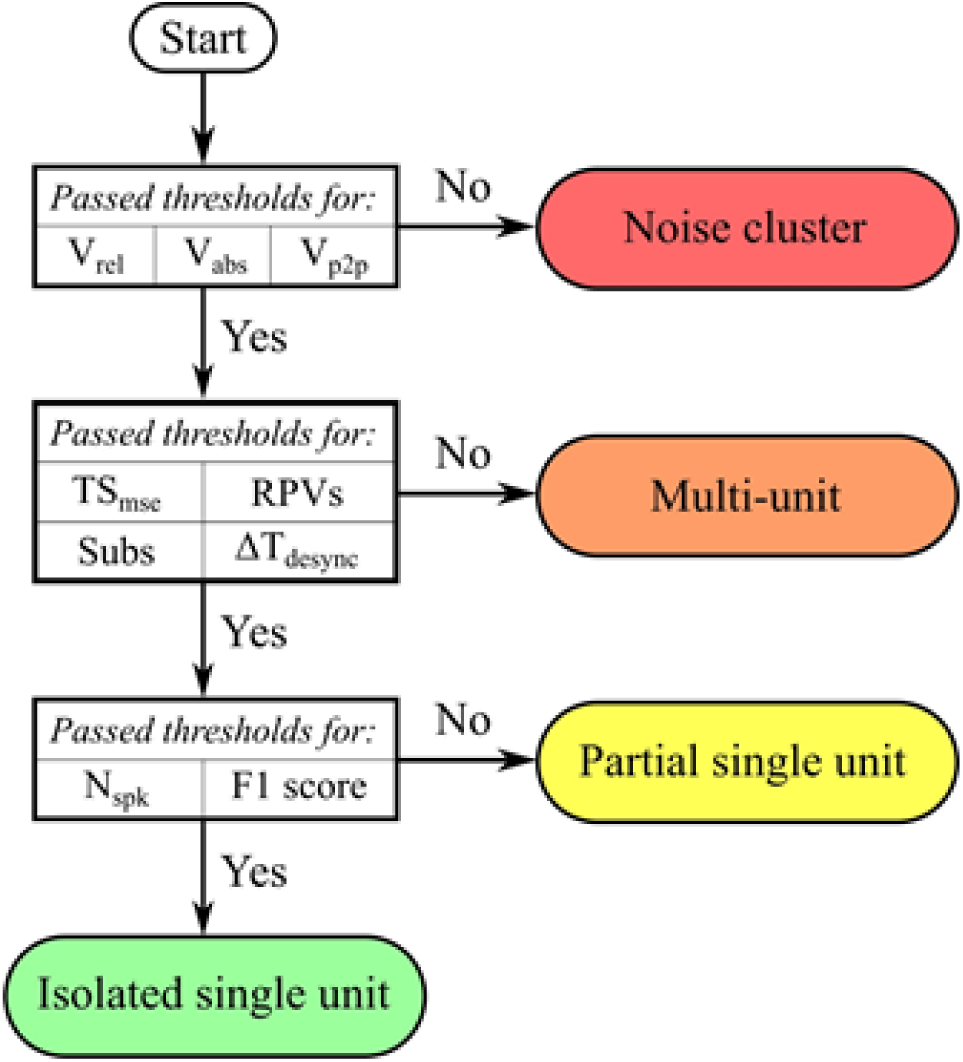
Flow-chart for determining the level of quality of a unit. See Table 1 for the description of each metric.

#### Step 3.10: Spike analysis

As with the LFP analysis (see Step 2.5), we also use FieldTrip to analyse and visualize the spiking data. PASER provides wrapper functions for plotting figures such as the inter-spike interval histogram, (joint) peri-stimulus time histogram, (cross-)correlogram and spike raster plot. To quantify differences between spike trains, PASER has a wrapper function for the external toolbox cSPIKE [42] to calculate spike train distance measures such as the (rate independent) SPIKE-distance [43, 44]. Lastly, an implementation of the multi-scale relevance to determine how relevant a neuron is for encoding the animal’s observed behaviour [45] is also included in the PASER toolbox.

### 2.4 Spike sorting

Spike sorting plays a central role in the data processing pipeline, because accurate identification of response properties of single neurons is essential for understanding the principles of neural coding and activity dependent expression of plasticity in networks [46–50]. A reason why response properties of single neurons cannot be predicted by the local population dynamics is that individual neurons can display significantly different firing patterns compared to adjacent neurons. For example, sparsely firing neurons can be tuned for particular stimuli and this selectively will be overlooked if their responses are not separated from the collective activity that is being measured [31, 38], since their sparse coding is often outweighed by the high firing rate of less selective neurons [51]. Therefore, we wish to assign each detected spike to the neuron that fired it, which is made possible because each neuron close to the electrode (< 50 µm [52]) fires spikes that appear in the signal as a uniquely distinctive waveform. This is due to the position of the neuron relative to the multiple contacts of the electrode, the cell’s morphology and the filtering properties of the medium [23,24,30,40,53,54]. Over the years many spike sorting algorithms have been developed to perform spike sorting independent from the differences in electro-physiological recording methods and hardware [55], complex background noise, highly non-Gaussian variance in waveform shape, electrode drift and time-overlapping spikes [53,56,57]. Therefore, PASER implements several of these methods. To determine the performance of spike sorting using each of these approaches, here we present a detailed performance evaluation.

#### 2.4.1 Method comparison

An ideal way of comparing the performance of spike sorting algorithms is through the use of ground-truth test data [52, 55], which is obtained from simultaneous intra- and extracellular recordings [31], resulting in a data set where the neuronal source of each spike is known a priori. However, such data sets are difficult to acquire and typically yield only a small number of simultaneously recorded neurons. Instead, we will investigate the efficiency of each method by running it on authentic in-vivo data [2] and compare spike sorting methods against each other based on computational speed, unit yield/quality and reliability (see below).

We limit ourselves to spike sorting methods that use the MATLAB language, since PASER as well as a majority of state-of-the-art toolboxes specialized in processing and analysis of extracellular recordings are developed in MATLAB [58]. This allows us to create a single-language toolbox, which improves its ease-of-use and development. Therefore, we exclude numerous open-source spike sorting programs written in other programming languages (primarily Python and C++), which pertains to recently published algorithms such as BOTMpy [39, 59], Combinato [33], Herding Spikes [60], Klusta [61], MountainSort [56], Spyking CIRCUS [41] and YASS [62]. As a result we evaluated nine different spike sorting algorithms, including CBPSpikesort [53], FMMSpikeSorter [63], ISO-SPLIT [64], KFMM [65], Kilosort [21], MClust [66], opass [67], OSort [68], UltraMegaSort2000 [38], WaveClus [69]. Note that the MClust and WaveClus methods both rely on super-paramagnetic clustering (SPC) and are therefore treated as a single method for this comparison. The SPC implementation adopted by the MClust toolbox is used in the analysis. Furthermore, the UltraMegaSort2000 algorithm has been adapted by vectorizing large sections of the original source code in order to speed up processing time by one order of magnitude. Since we are only interested in the final output of each algorithm, we will not go into details about their modes of operation, but instead refer the reader to the respective papers for further details. Treating the methods as black boxes is also key for keeping the PASER toolbox modular and plugin-based. Not tying the toolbox to any particular spike sorting routine makes it future-proof, because it will be easier to incorporate a new and improved method when it is released later down the road.

To carry out the comparison, we run the aforementioned spike sorting algorithms using default parameters on extracellular recordings obtained in freely behaving mice using measured by tetrodes, using a 64-bit Windows 10 PC with a quad-core 3.40GHz CPU, GeForce GTX 1060 6GB GPU and 16GB of RAM. Where possible, sub-routines are shared across different spike sorting methods, such as spike detection and feature extraction, in order to bring about a fairer comparison. The ISO, KFM and SPC methods require a lower dimensional representation of spike waveforms as input to their respective pipelines. In all three cases, we extract 12-dimensional feature vectors for every waveform using principal component analysis, where the feature vectors are constructed by taking the first three principal components of each waveform channel and then concatenating the feature values of the four channels [70]. Furthermore, FMM, ISO, KFM, OPS, OST, SPC and UMS all necessitate spike detection, which is performed using a threshold of 3 MAD. CBP and KST, on the other hand, are template matching approaches that operate directly on the filtered time series data, which assume that the signal can be decomposed as a sum of templates plus noise [71].

#### 2.4.2 Computational time

If we run a neural recording session for an hour with 16 tetrodes simultaneously sampling at a rate of 30 kHz, then we have to process several billions of data points, containing hundreds of thousands of spikes in total. When dealing with such large datasets, we have to make sure that the data can be processed within a realistic time frame, e.g. no longer than an order of magnitude beyond the time it took to record the data. Spike sorting methods that are poorly optimized in this respect must immediately be disregarded, even if they deliver accurate results. For this reason, we first evaluate the duration of spike sorting for the selected methods for data sections of 100 and 1000 seconds, where each duration is repeated 16 times (Figure 8). The CBP, KFM and OPS algorithms exceeded the maximum computational time of 10 minutes that was allotted for this analysis and are therefore excluded from further testing. The remaining six methods show similar computational speeds for the 100 seconds data section, but start to display significantly different results when scaled to 1000 seconds. For FMM and KST, the computational time scales well to the larger data set, but for OST and UMS the scaling is poor.

**Figure 8:**
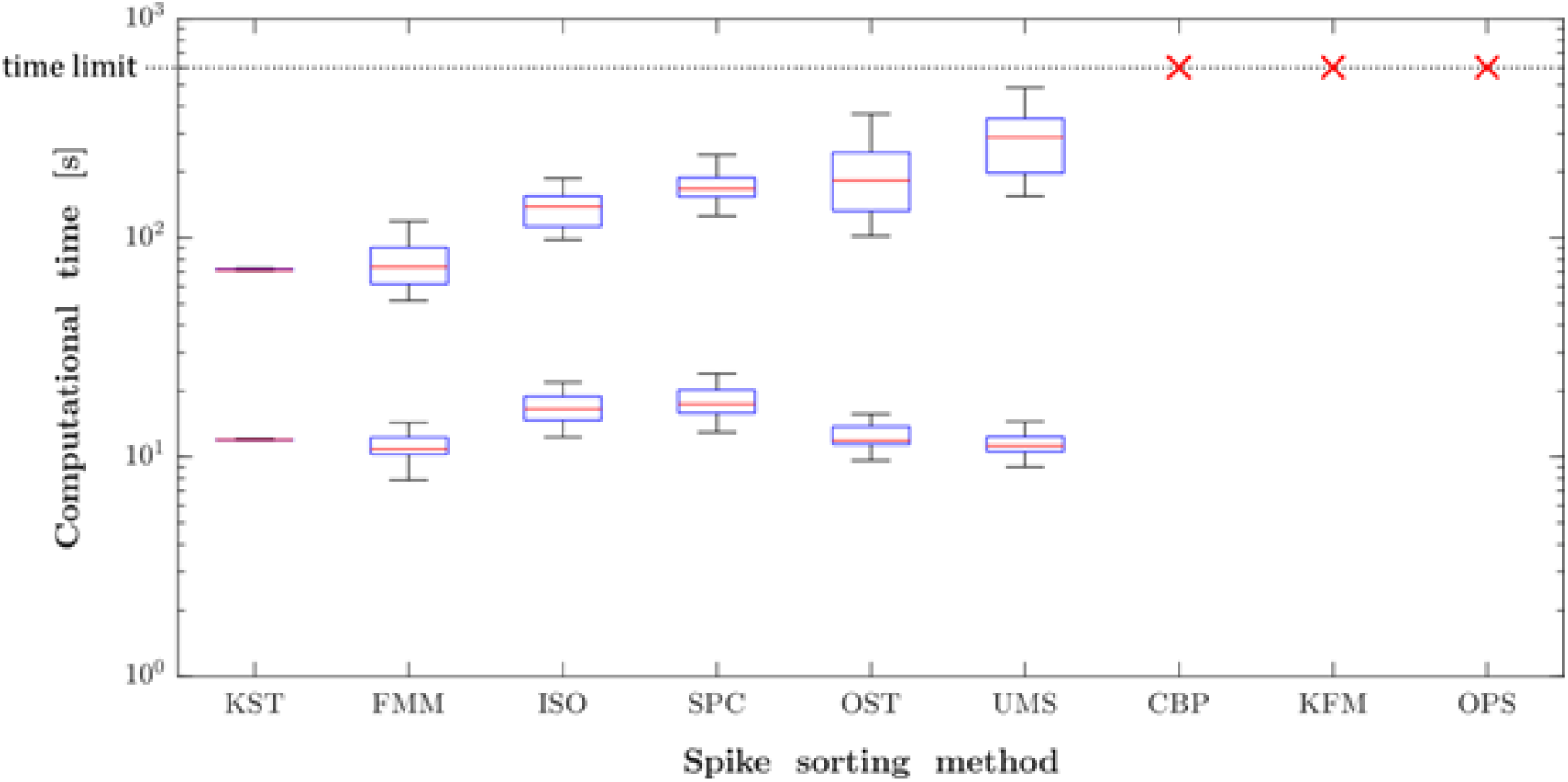
Computational efficacy of spike sorting algorithms. The lower and upper boxes respectively indicate the timing results from processing data sections of 100 and 1000 sec in duration. The spike sorting methods compared are CBPSpikesort (CBP), FMMSpikeSorter (FMM), ISO-SPLIT(ISO), KFMM (KFM), Kilosort (KST), MClust (SCS), opass (OPS), OSort (OST), UltraMegaSort2000 (UMS), WaveClus (SCS). The time limit of 10 minutes was exceeded by the CBP, KFM and OPS methods.

#### 2.4.3 Unit yield and quality

Figure 9 shows how many units of each quality (see Figure 7) have been obtained for the six remaining methods for 1000 seconds of data recorded by 16 tetrodes. The relative unit isolation performance of ISO, OST and UMS cannot be justified given their computational speed, so these methods are rejected from the rest of the analysis. The remaining three approaches are implemented on three more recording sessions of identical duration, but with varying degrees of noise. The results of this investigation are shown in Figure 10. In contrast to the first case, the unit yield of SPC is now drastically smaller compared to FMM and KST. For this reason, only FMM and KST are considered for the last assessment.

**Figure 9:**
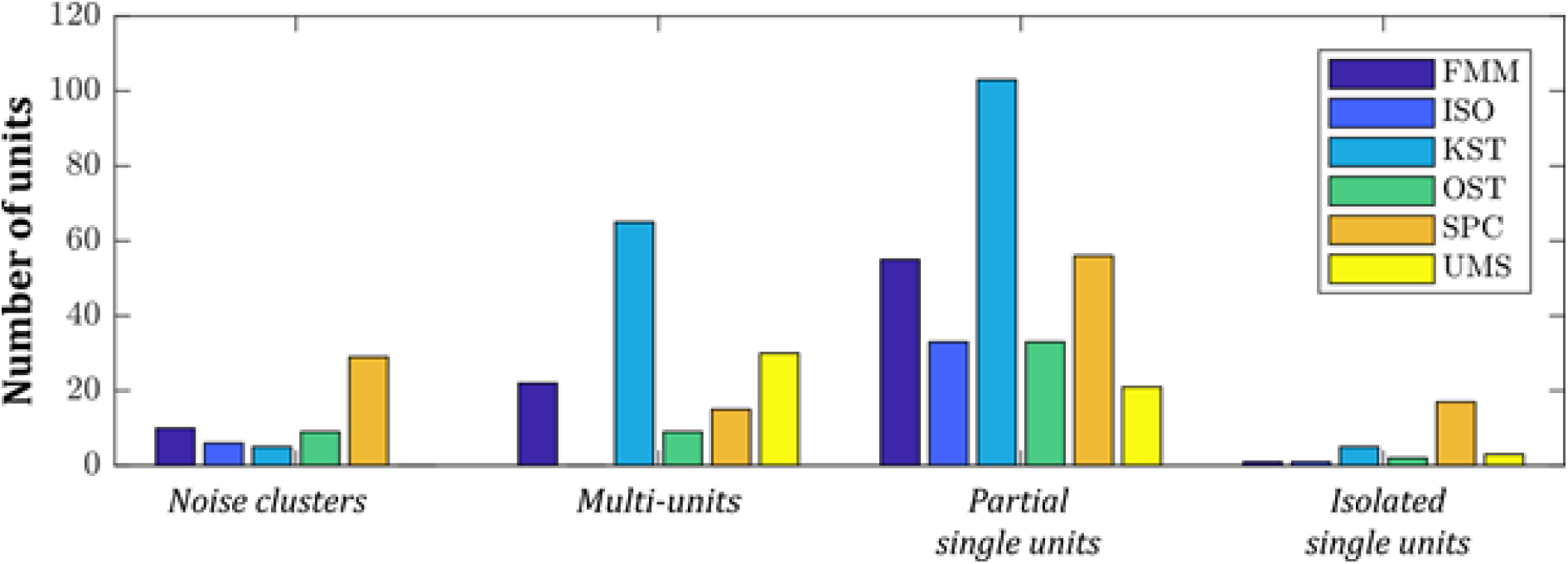
Unit yield and quality. For every unit quality type (noise cluster, multi-unit, partial and isolated single unit), a histogram is shown with the number of units obtained for that type by each of the six spike sorting methods (FMM, ISO, KST, OST, SPC and UMS). Each method processed the same data set of 1000 seconds recorded by 16 tetrodes.

**Figure 10:**
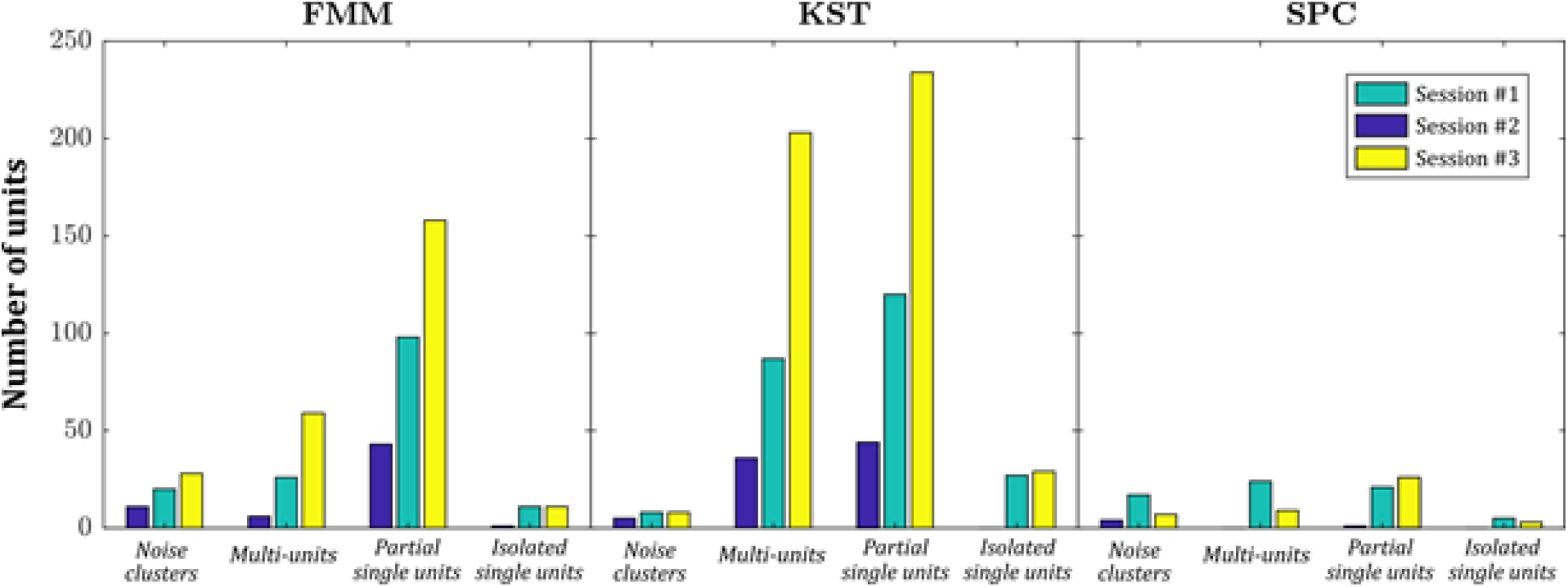
As in Figure 9 but now for three different recording sessions of 1000 seconds using 16 tetrodes and only for the three remaining spike sorting methods: FMM (left), KST (centre) and SPC (right).

#### 2.4.4 Reliability

Spike sorting results are more reliable when the output units are stable across multiple repetitions, so for the final evaluation we test the degree of repeatability of the remaining two methods. Of course, when the spike sorting method is deterministic, the output results will be identical if applied on the same data set, so we have to slightly perturb the data for each repetition in order to obtain a meaningful measure of stability. Two different perturbation techniques are employed, which are *self-blurring* and *noise-reversal* [29]. In the former case, we calculate for every spike the difference between its waveform and the mean waveform of its cluster, which results in a noise distribution for that cluster. A random permutation of the noise distribution is then added to every spike in the cluster, where we scale the addition by 0.3. This has the effect of broadening (blurring) the noise range of the cluster, but does not change the general waveform shape. Noise-reversal, on the other hand, works by flipping the sign of the background noise and reflecting the spike noise around the mean waveform of the cluster. So, for example, if a particular spike waveform has an amplitude that is 10 *µV* larger than the mean waveform, after noise-reversal it will be 10 *µV* smaller. After applying these perturbations, we check the correspondence of the results between the unperturbed, self-blurred and noise-reversed data by calculating a normalized confusion matrix. Each spike is put in a cell of the confusion matrix depending on the cluster it was assigned to in the first and second data set (e.g. unperturbed and self-blurred). Our stability measure is defined as the average value of the main diagonal of the confusion matrix. For FMM and KST, we respectively obtain average stability measures of 0.71*±*0.16 and 0.61*±*0.22 when considering partial and isolated single units, where the error is given by the standard deviation.

#### 2.4.5 The choice of spike sorting algorithm

We have evaluated nine different open-source spike sorting algorithms on the basis of computational speed, unit yield/quality and reliability (stability) using in-vivo extracellular tetrode recordings in freely behaving mice [2]. Two spike sorting algorithms demonstrated significantly superior results over the other methods. These are the dictionary learning and mixture modelling approach of FMM, together with the template matching plus GPU acceleration technique of KST. When contrasting these two methods, we observed approximately equal processing time for 1000 second data. KST, however, retrieved a greater number of high-quality units for three separate recordings relative to FMM. Conversely, FMM displayed greater stability of results over multiple repetitions. Ultimately, we opted to use KST as the default method for the PASER toolbox, where the deciding factor is that KST scales better to very long recording sessions and to higher-density probes with respect to computational speed (data not shown). This choice does not mean that the performance of KST is universally and objectively better than all other investigated methods, but rather that KST superiorly fits our subjective needs of ease-of-use, efficiency and quality of results for our particular set-up. PASER user can opt to integrate any of the implemented options, although the default of the toolbox is the KST.

### 2.5 Data visualization and quality control

To evaluate the fidelity of the spike sorting output for each identified unit on an individual basis PASER provides tools for automatic quality assessment as well as visualization schemes for manual curation, some of which were already introduced in Section 2.3 and will be further expanded in this section.

#### 2.5.1 Visualization

Figure 11 is a representation of the typical visualization that PASER generates for every cluster, which the user can use to manually judge the quality of the cluster, to confirm the automated classification of the units as described in Step 3.9.

**Figure 11.**
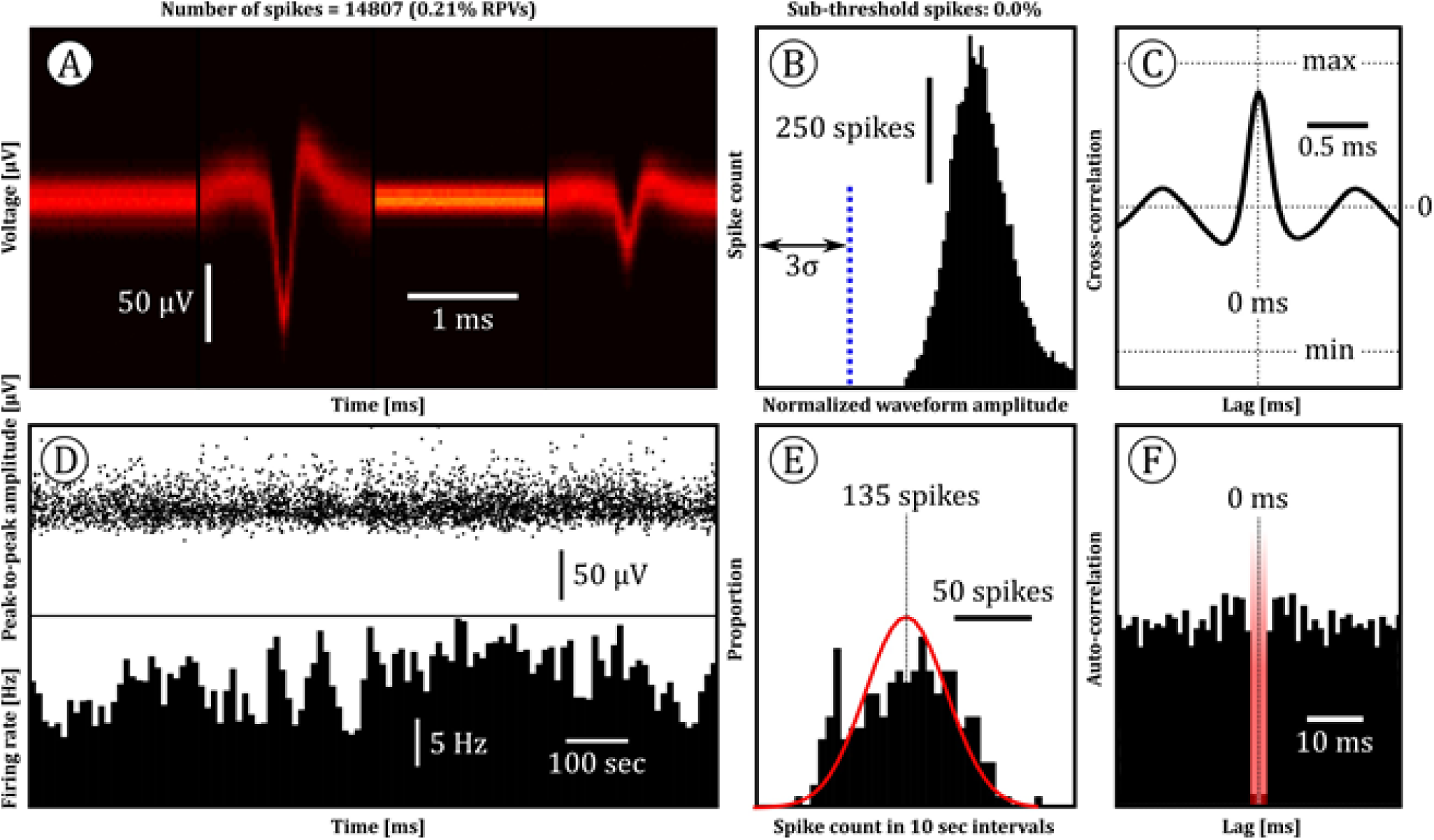
Default quality metrics displayed for each cluster. (A) Concatenated spike shapes across the tetrode channels visualized as a density heatmap of time-voltage values. Total number of detected spikes and the proportion of spikes that violate the refractory period (RPV) are noted in the figurine title. (B) Histogram of normalized spike amplitudes. Any amplitude to the right of the dotted vertical line crosses the minimum threshold for spike detection, which is three times the background noise (σ). (C) Cross-correlation between the two tetrode channels that are maximally dissimilar. The large peak at zero lag indicates that the spike waveforms are temporally aligned. (D) The peak-to-peak amplitude (top) and firing rate (bottom) over the whole recording period in a session, showing the temporal stability of the cluster. (E) Distribution of running average spike counts in 10 sec bins (with 5 sec overlap). The red curve indicates the predicted distribution of spike counts by a Gaussian distribution given the observed variance in spike count. (F) Autocorrelation of spike events. The red bar in the centre of the plot marks the refractory period.

#### 2.5.2 Amplitude

At the very least, a high-quality cluster should have a mean waveform amplitude that is significantly greater than the default spike detection threshold of 3 times the background noise level, σ. This is a requirement, because the default spike threshold is purposefully set to an exceedingly low level in order to not miss any spikes, but the median waveform of a genuine single unit should have a better signal-to-noise ratio, which is why the minimum median amplitude is set at 4σ by default. Furthermore, biologically unrealistic excessively high amplitudes are not acceptable either, as these are indicative of artifacts or possibly axonal signals (see Table 1 for amplitude threshold values).

Figure 11(B) shows a histogram of the spike waveform amplitudes in the cluster. Following Hill et al. (2011), we can use this distribution of amplitudes to estimate the number of spikes that does not cross the detection threshold if we assume that the amplitudes are distributed according to a distribution determined a priori (e.g. a Gaussian distribution) [38]. Note that this method is only sensible when actually using a spike detection threshold, which does not apply to template matching techniques, like the KiloSort (KST), where we can directly calculate the fraction of sub-threshold spikes in the cluster. For all other cases, Figure 12 shows a representative visualization of how the fraction of sub-threshold spikes is computed. The plot shows a histogram of normalized amplitude values, which are obtained by dividing the absolute amplitudes by the spike detection threshold, set to 3σ. We then fit a Gaussian on this distribution, while taking into consideration that the left tail might be cut off. The region below the fitted curve and to the left of the spike detection threshold then corresponds with the fraction of sub-threshold spikes (orange shaded area). In our quality control scheme, single units can have up to a maximum of 10% of their spikes below the detection threshold.

**Figure 12:**
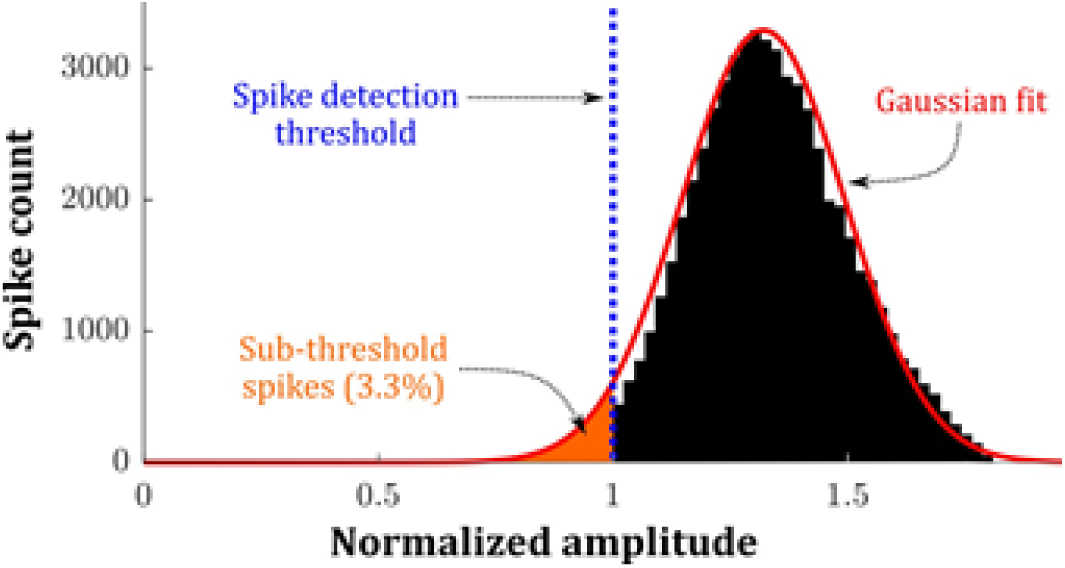
Process to estimate the percentage of sub-threshold spikes in a unit. The black bar plot is a histogram of normalized waveform amplitudes of a representative unit. The amplitudes are normalized through dividing by the spike detection threshold (3σ), indicated by the dotted vertical line. A Gaussian is fitted on the amplitude distribution, where the percentage of sub-threshold spikes is estimated by the (orange) area under the Gaussian curve to the left of the spike detection threshold divided by the total area under the Gaussian.

#### 2.5.3 Firing rate stability

Non-stationarities in the amplitude is often evidence for electrode drift or tissue movement, while atypical deviations in firing rate are indicative of noise contamination. The latter is quantified by segmenting the recording session into 10 second intervals, with 5 second overlap, and counting the number of spikes that occur in each segment. Empirically, we have determined that this distribution of spike counts typically follow a Gaussian distribution with a standard deviation of λ^3/2^, where λ is the average number of spikes per interval. In Figure 11(E), this theoretical distribution is shown as a red curve, where λ = 135, while the histogram of spike counts is shown in black. As a single evaluation metric for the firing rate stability of the unit, we use the ratio of the dispersion around λ between the theoretical and observed spike count distributions. Ideally, this ratio should be 1, but we allow the observed distribution to have a dispersion value that is at most 4 times greater than the dispersion of the theoretical distribution. For our measure of dispersion, we use the average MSE between λ and each spike count:

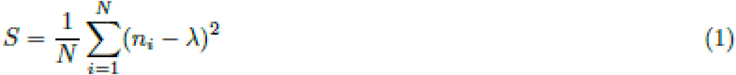

Where S is the dispersion, N is the total number of 10 sec intervals in the session and ni is the spike count in interval i. The firing rate stability measure, F Smse, is then calculated through:

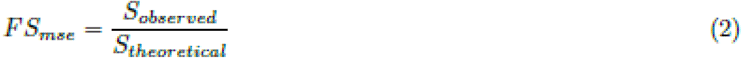

The benefit of using the MSE is that extreme outliers in the observed spike count distribution incur a heavy penalty to the stability measure, even when we only have a few of them, since we are squaring the difference. This is preferable because these types of outliers are never conventional and should significantly degrade the quality of the unit.

In Figure 13, quality control plots are shown for a unit with highly non-stationary firing rates. The spikes of the unit appear in bursts with limited activity in between. Such spiking behaviour is typically due to noise corruption, where the bursting sections coincide with noisy regions in the raw data. Unsurprisingly, the theoretical and observed spike count distributions are significantly divergent (see bottom-right plot in Figure 13). The theoretical distribution has many more occurrences of small and large spike counts compared to what is expected from a stable unit, resulting from the silent and bursting sections. For this unit we obtain a FSmse of 12.6, which exceeds the maximum allowable value, so we do not classify it as a high-quality single unit (as per the criteria detailed in Figure 7). In contrast, the FSmse of the unit in Figure 11 is 1.5. This disparity in quality between the two units would not have been elucidated if we had judged them by their waveform shape alone.

**Figure 13:**
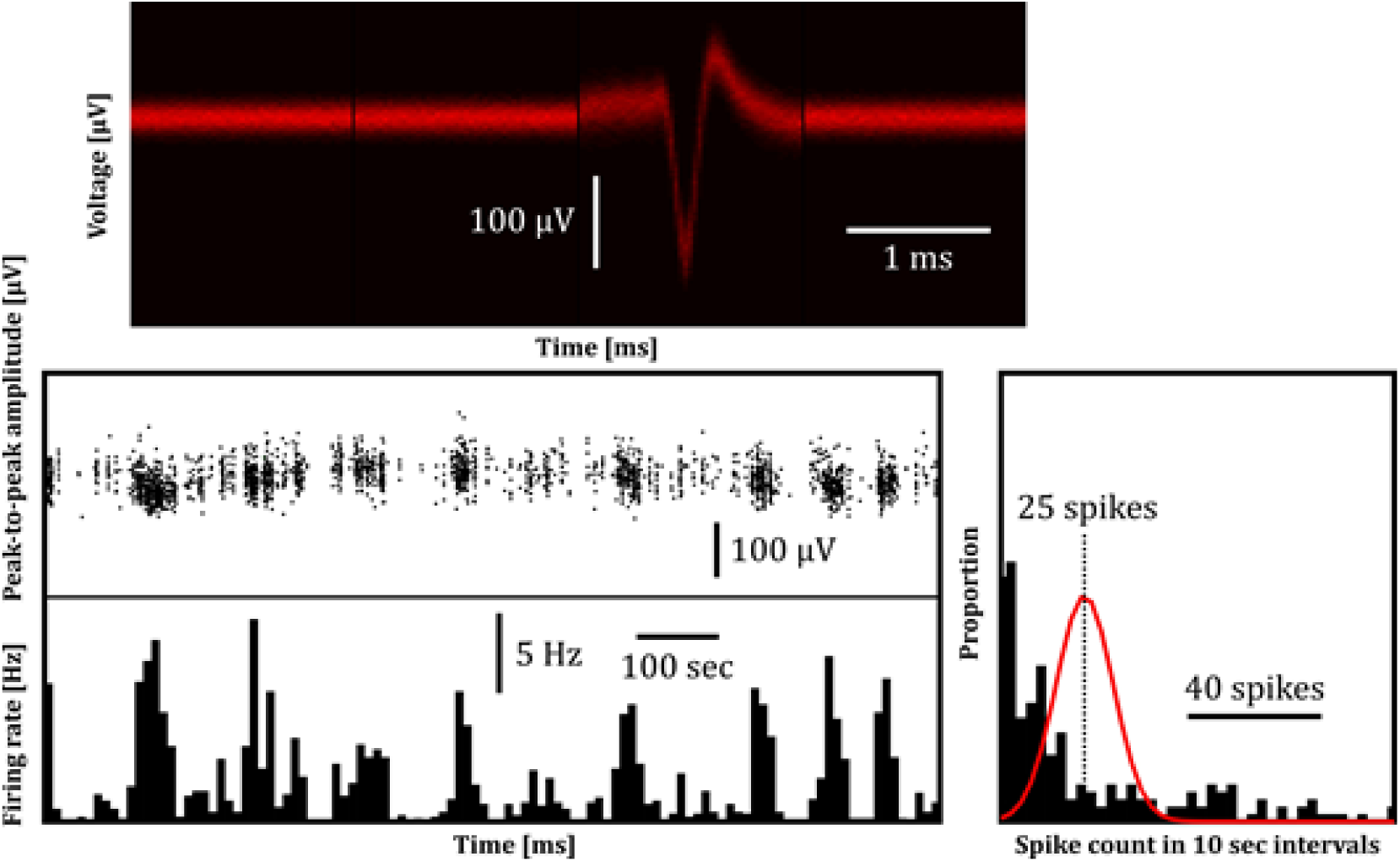
Example of quality control plots for a unit with poor firing rate stability. See Figure 11 for descriptions of the three sub-plot types. The highly irregular firing pattern potentially results from systematic noise influencing the waveform shape, leading to a sub-cluster that only shows activity whenever the noise is present.

Further stability analyses are also possible, such as carrying out linear regression on the waveform amplitudes as a function of time (see top-plot in Figure 11(D)), where a regression slope that strongly deviates from zero would be an indication of non-stationarity. Additionally, the fidelity of the Gaussian fit on the empirical amplitude distribution (see Figure 12), where a poor fit also suggests irregular amplitudes, can be visualized.

#### 2.5.4 Refractory period

Following an action potential, there exists a short period of time where a neuron cannot generate another action potential, termed the refractory period. Whenever we evaluate the quality of a unit, we should check how many of its spikes occur within the refractory period (set to 1.5 ms) of the previous spike in the cluster, known as refractory period violations (RPVs). These RPVs can be caused either by noise contamination or the superimposed activity of another neuron. In the latter case, we are dealing with a multi-unit and we should classify the cluster accordingly. The frequency of RPVs can also be inferred from Figure 11(F), which shows the autocorrelation of the clustered spikes after binning them into 1 ms bins. A significant dip in the autocorrelation function is expected around zero lag, up to the refractory period (marked by the red shaded area).

#### 2.5.5 Isolation quality

Lastly, we require a means of quantifying the degree to which a cluster uniquely represent the activity from a single neuron. This measure is generally referred to as the isolation quality of a unit, which is typically gauged by looking at differences in waveform shapes between clusters measured on the same polytrode. Various methods to assess this difference have been identified in the literature, which all necessitate that the spike sorted waveforms are represented in a lower dimensional feature space, using a dimensionality reduction technique such as principal component analysis (PCA). This procedure is depicted in Figure 14, where the high dimensional waveforms are reduced to their first two principal components after performing PCA, resulting in two easily visualized clusters in two-dimensional feature space. To determine how isolated each of these clusters are, we can use measures such as the Lratio [72] or the Isolation Distance [70], both of which employ the Mahalanobis distance to quantify the level of separation [73]. Rather than exploring the exact details of each metric, we instead note that both methods come with drawbacks. The Isolation Distance is ill-defined for large clusters [73], while the Lratio is difficult to interpret [74]. Furthermore, both methods lack a global scale, which makes it infeasible to set a general threshold for differentiating between well and poorly isolated units [75].

**Figure 14:**
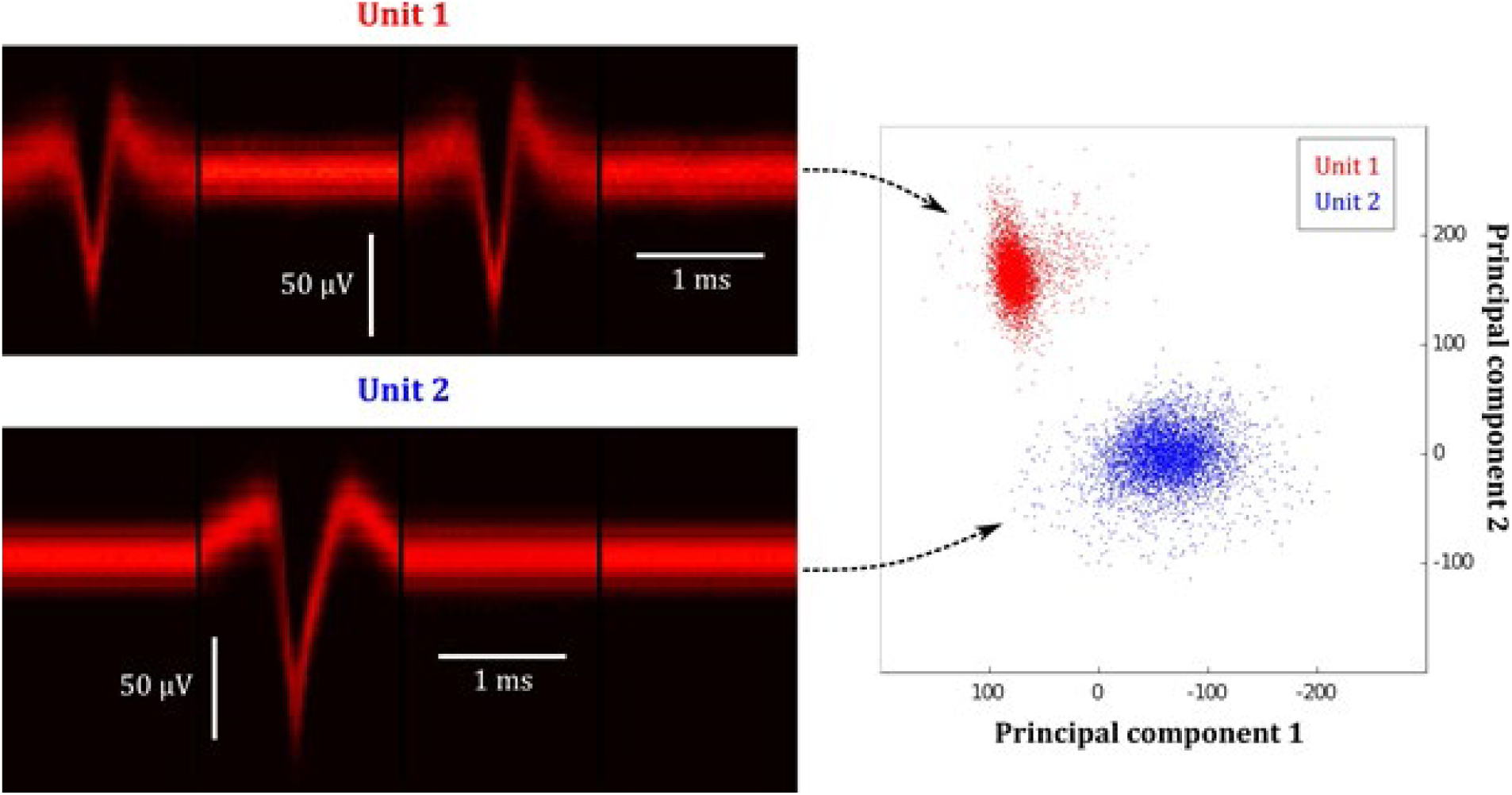
Separation distance between units. To determine how well each unit is isolated from neighboring clusters, we reduce the dimensionality of their corresponding waveforms using PCA and calculate inter-cluster distances in feature space. In the scatter plot on the right, each dot indicates the first two principal component values for an individual spike in the corresponding unit.

As an alternative, we can go with a model-based approach, where we assume that the clusters in feature space can be modelled according to a particular distribution and calculate isolation quality metrics based on that assumption. For example, we can fit a multivariate Gaussian mixture model on the clustered data and then compute the probability of false-positive (FP) and false-negative (FN) errors for each unit, under the assumption of a Gaussian cluster shape [38, 60]. A particular unit has FP errors when spikes are misassigned to a different cluster. In contrast, FN errors occur when spikes associated with the unit were missed, discarded or grouped with some other unit [29]. The downside of using a mixture of Gaussians to calculate FP and FN errors is that it does not adequately account for cluster drift and heavy tails observed in experimental data [74]. For this reason, we incorporate the method developed by Shan et al. (2017), whereby a mixture of drifting t-distributions is fitted on the clustered data, which gives increased performance over a mixture of Gaussians for long chronic recordings [74]. However, instead of letting the resulting FP and FN probabilities represent the final isolation quality of a cluster, we condense both values into a single evaluation metric, the F1-score, to simplify quality classification:

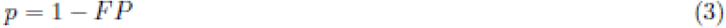

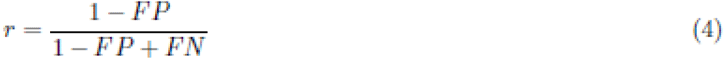

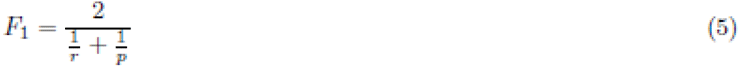

Where p and r are precision and recall respectively. The harmonic mean between p and r then evaluates to the F1-score, where we uphold the criterion that isolated single units should have F1 > 0.9, which is passed by both units in Figure 14. In some cases, it might be wiser to consider FP and FN errors separately since they have different implications. Random FN errors can lead to an underestimation of firing rate, while FP errors can have the more disastrous consequence of wrong conclusions being drawn about information encoding [52], so we may want to exercise greater caution when confronted with FPs.

## 3 Discussion

We have presented a methodology for processing chronic extracellular neural recordings, where existing methods from the literature and self-developed techniques are combined to achieve an acceptable trade-off between automation and speed while maximizing the accuracy in signal processing. The resulting processing pipeline has been integrated into a MATLAB toolbox named PASER, which is available for download at github.com/DepartmentofNeurophysiology/Paser. We paid particular attention to the detection, categorization and refinement of action potentials identified in the voltage trace measured by the intracranial multichannel electrodes.

A key step in this routine is the spike sorting process: the assignment of spikes to the source neuron that fired it. A wide variety of open-source spike sorting algorithms are freely available online, with varying levels of performance. Currently, no spike sorting algorithm exists that consistently demonstrates a high degree of accuracy across a broad range of experimental conditions. This is partly due to a lack of validation methods, since ground truth data is difficult to come by. A potential solution to this problem is measuring the reliability of spike sorting algorithms on synthetic data, which can be generated using complex biophysical simulations or by extracting isolated spikes from real data and then randomly inserting them into a different recording [52, 55]. However, reproducing important features of real recordings, e.g. non-Gaussian clusters and multi-unit activity [31], remains a serious modelling challenge.

Additionally, issues such as overlapping spikes and nonstationarities, which complicate the spike sorting process, continue to be largely unresolved. Spike overlap occurs whenever two nearby neurons fire in short succession of each other, leading to a complex superposition of their two waveforms in the electrode signal. This waveform superposition is often significantly distinct from the waveforms of the two constituent neurons, which means that the spikes are either discarded or put into an entirely separate cluster [76]. Template matching techniques can provide a partially automated solution to deal with overlapping spikes [57]. Furthermore, long lasting chronic recordings in-vivo introduce the issue of non-stationary spike waveforms, resulting from electrode drift, tissue relaxation, bursting activity, prolonged firing, changing cellular factors and dynamic experimental protocols [52,53,65,71], which can gradually alter both the amplitude and shape of the neuron’s waveform over time. A basic, but non-ideal, way of accounting for nonstationarities is to spike sort the data in temporally segmented blocks and then merging clusters that belong to the same unit across these blocks [33, 77].

However, more robust frameworks are required when carrying out spike sorting in real-time [71]. Among other reasons, real-time spike sorting is attractive due to the substantial reduction in data transmission and storage [68], which becomes especially advantageous when recording with high-density probes for long sessions at the typically sampling rate of extracellular signals (around 30 kHz). The massive size of the subsequent raw voltage traces will make it infeasible to store them for offline processing. Moreover, real-time spike sorting allows for experimental conditions that adapt according to the neural responses that are observed which makes room for wider scientific investigation including but not limited to the neural basis of adaptive sensorimotor computation [78, 79], contextual information processing [80–83] and storage [84–87], navigation [88–90], and information transfer in neural circuits [46, 48]. Brain-machine-interfaces (BMI), like limb prosthetics, also necessitate that spike sorting is performed in real-time on a time-scale of hundreds of milliseconds [65], as these are usually controlled by direct neuronal signalling that is measured invasively by an array of electrodes [91]. Although, for BMI applications, the exact identity of each spike may actually not be crucial and the spike sorting stage can be omitted entirely [52].

Looking towards the future, it is imperative that the presented toolbox remains under active development, so that it can be rapidly updated whenever novel methodology is published that demonstrates improved performance in any of the aforementioned areas or when state-of-the art algorithms are released that are able to fully resolve some of the outstanding problems (e.g. overlapping spikes). For this reason, PASER is made modular and has not been integrated into a graphical user interface, which may increase user-friendliness, but can also constrain development. Ultimately, we wish to end up with a data processing pipeline that is able to detect and sort spikes with higher speed and accuracy, as well as more objectively assess the quality of the resulting units, while maintaining full automation.

## References

[1] Leibiger I B and Berggren P-O 2015 Regulation of glucose homeostasis using radiogenetics and magnetogenetics in mice. Nat. Med. 21 14–6

[2] Kole K, Zhang Y, Jansen E J R, Brouns T, Biljsma A, Calcini N, Yan X, Lantyer A da S and Celikel T Assessing the utility of MAGNETO to control neuronal excitability in the somatosensory cortex Nat Neurosci, in press

[3] Barker A T, Jalinous R and Freeston I L 1985 Non-invasive magnetic stimulation of human motor cortex. Lancet 1 1106–7

[4] Tufail Y, Matyushov A, Baldwin N, Tauchmann M L, Georges J, Yoshihiro A, Tillery S I H and Tyler W J 2010 Transcranial pulsed ultrasound stimulates intact brain circuits. Neuron 66 681–94

[5] Etoc F, Vicario C, Lisse D, Siaugue J-M, Piehler J, Coppey M and Dahan M 2015 Magnetogenetic control of protein gradients inside living cells with high spatial and temporal resolution. Nano Lett. 15 3487–94

[6] Maggioni E, Arrubla J, Warbrick T, Dammers J, Bianchi A M, Reni G, Tosetti M, Neuner I and Shah N J 2014 Removal of pulse artefact from EEG data recorded in MR environment at 3T. Setting of ICA parameters for marking artefactual components: application to resting-state data. PLoS ONE 9 e112147

[7] Mullinger K J, Havenhand J and Bowtell R 2013 Identifying the sources of the pulse artefact in EEG recordings made inside an MR scanner. Neuroimage 71 75–83

[8] Allen P J, Josephs O and Turner R 2000 A method for removing imaging artifact from continuous EEG recorded during functional MRI. Neuroimage 12 230–9

[9] Mantini D, Perrucci M G, Cugini S, Ferretti A, Romani G L and Del Gratta C 2007 Complete artifact removal for EEG recorded during continuous fMRI using independent component analysis. Neuroimage 34 598–607

[10] Jung T P, Makeig S, Humphries C, Lee T W, McKeown M J, Iragui V and Sejnowski T J 2000 Removing electroencephalographic artifacts by blind source separation. Psychophysiology 37 163–78

[11] Gordon S M, Lawhern V, Passaro A D and McDowell K 2015 Informed decomposition of electroencephalographic data. J. Neurosci. Methods 256 41–55

[12] Plöchl M, Ossandón J P and König P 2012 Combining EEG and eye tracking: identification, characterization, and correction of eye movement artifacts in electroencephalographic data. Front. Hum. Neurosci. 6 278

[13] Srivastava G, Crottaz-Herbette S, Lau K M, Glover G H and Menon V 2005 ICA-based procedures for removing ballistocardiogram artifacts from EEG data acquired in the MRI scanner. Neuroimage 24 50–60

[14] Briselli E, Garreffa G, Bianchi L, Bianciardi M, Macaluso E, Abbafati M, Grazia Marciani M and Maraviglia B 2006 An independent component analysis-based approach on ballistocardiogram artifact removing. Magn. Reson. Imaging 24 393–400

[15] Mantini D, Perrucci M G, Del Gratta C, Romani G L and Corbetta M 2007 Electrophysiological signatures of resting state networks in the human brain. Proc Natl Acad Sci USA 104 13170–5

[16] Neuner I, Arrubla J, Felder J and Shah N J 2014 Simultaneous EEG-fMRI acquisition at low, high and ultra-high magnetic fields up to 9.4 T: perspectives and challenges. Neuroimage 102 Pt 1 71–9

[17] Neuner I, Warbrick T, Arrubla J, Felder J, Celik A, Reske M, Boers F and Shah N J 2013 EEG acquisition in ultra-high static magnetic fields up to 9.4 T. Neuroimage 68 214–20

[18] Debener S, Strobel A, Sorger B, Peters J, Kranczioch C, Engel A K and Goebel R 2007 Improved quality of auditory event-related potentials recorded simultaneously with 3-T fMRI: removal of the ballistocardiogram artefact. Neuroimage 34 587–97

[19] Vanderperren K, De Vos M, Ramautar J R, Novitskiy N, Mennes M, Assecondi S, Vanrumste B, Stiers P, Van den Bergh B R H, Wagemans J, Lagae L, Sunaert S and Van Huffel S 2010 Removal of BCG artifacts from EEG recordings inside the MR scanner: a comparison of methodological and validation-related aspects. Neuroimage 50 920–34

[20] Siegle J H, López A C, Patel Y A, Abramov K, Ohayon S and Voigts J 2017 Open Ephys: an open-source, plugin-based platform for multichannel electrophysiology. J. Neural Eng. 14 045003

[21] Pachitariu M, Steinmetz N, Kadir S, Carandini M and Harris K D 2016 Kilosort: realtime spike-sorting for extracellular electrophysiology with hundreds of channels BioRxiv

[22] Oostenveld R, Fries P, Maris E and Schoffelen J-M 2011 FieldTrip: Open source software for advanced analysis of MEG, EEG, and invasive electrophysiological data. Comput. Intell. Neurosci. 2011 156869

[23] Allen C B, Celikel T and Feldman D E 2003 Long-term depression induced by sensory deprivation during cortical map plasticity in vivo. Nat. Neurosci. 6 291–9

[24] Celikel T, Szostak V A and Feldman D E 2004 Modulation of spike timing by sensory deprivation during induction of cortical map plasticity. Nat. Neurosci. 7 534–41

[25] Foeller E, Celikel T and Feldman D E 2005 Inhibitory sharpening of receptive fields contributes to whisker map plasticity in rat somatosensory cortex. J. Neurophysiol. 94 4387–400

[26] Voigts J, Siegle J H, Pritchett D L and Moore C I 2013 The flexDrive: an ultra-light implant for optical control and highly parallel chronic recording of neuronal ensembles in freely moving mice. Front. Syst. Neurosci. 7 8

[27] Jia X, Siegle J H, Bennett C, Gale S D, Denman D J, Koch C and Olsen S R 2019 High-density extracellular probes reveal dendritic backpropagation and facilitate neuron classification. J. Neurophysiol. 121 1831–47

[28] Jun J J, Steinmetz N A, Siegle J H, Denman D J, Bauza M, Barbarits B, Lee A K, Anastassiou C A, Andrei A, Aydın Ç, Barbic M, Blanche T J, Bonin V, Couto J, Dutta B, Gratiy S L, Gutnisky D A, Häusser M, Karsh B, Ledochowitsch P and Harris T D 2017 Fully integrated silicon probes for high-density recording of neural activity. Nature 551 232–6

[29] Barnett A H, Magland J F and Greengard L F 2016 Validation of neural spike sorting algorithms without ground-truth information. J. Neurosci. Methods 264 65–77

[30] Jun J J, Mitelut C, Lai C, Gratiy S, Anastassiou C and Harris T D 2017 Real-time spike sorting platform for high-density extracellular probes with ground-truth validation and drift correction BioRxiv

[31] Rey H G, Pedreira C and Quian Quiroga R 2015 Past, present and future of spike sorting techniques. Brain Res. Bull. 119 106–17

[32] Quian Quiroga R 2009 What is the real shape of extracellular spikes? J. Neurosci. Methods 177 194–8

[33] Niediek J, Boström J, Elger C E and Mormann F 2016 Reliable Analysis of Single-Unit Recordings from the Human Brain under Noisy Conditions: Tracking Neurons over Hours. PLoS ONE 11 e0166598

[34] Ludwig K A, Miriani R M, Langhals N B, Joseph M D, Anderson D J and Kipke D R 2009 Using a common average reference to improve cortical neuron recordings from microelectrode arrays. J. Neurophysiol. 101 1679–89

[35] Kwon K Y, Eldawlatly S and Oweiss K 2012 NeuroQuest: a comprehensive analysis tool for extracellular neural ensemble recordings. J. Neurosci. Methods 204 189–201

[36] Bakštein E, Sieger T, Wild J, Novák D, Schneider J, Vostatek P, Urgošík D and Jech R 2017 Methods for automatic detection of artifacts in microelectrode recordings. J. Neurosci. Methods 290 39–51

[37] Dolan K, Martens H C F, Schuurman P R and Bour L J 2009 Automatic noise-level detection for extra-cellular micro-electrode recordings. Med. Biol. Eng. Comput. 47 791–800

[38] Hill D N, Mehta S B and Kleinfeld D 2011 Quality metrics to accompany spike sorting of extracellular signals. J. Neurosci. 31 8699–705

[39] Franke F, Quian Quiroga R, Hierlemann A and Obermayer K 2015 Bayes optimal template matching for spike sorting - combining fisher discriminant analysis with optimal filtering. J. Comput. Neurosci. 38 439–59

[40] Gao H, Solages C de and Lena C 2012 Tetrode recordings in the cerebellar cortex. J. Physiol. Paris 106 128–36

[41] Yger P, Spampinato G L, Esposito E, Lefebvre B, Deny S, Gardella C, Stimberg M, Jetter F, Zeck G, Picaud S, Duebel J and Marre O 2018 A spike sorting toolbox for up to thousands of electrodes validated with ground truth recordings in vitro and in vivo. elife 7

[42] Satuvuori E, Mulansky M, Bozanic N, Malvestio I, Zeldenrust F, Lenk K and Kreuz T 2017 Measures of spike train synchrony for data with multiple time scales. J. Neurosci. Methods 287 25–38

[43] Kreuz T, Chicharro D, Houghton C, Andrzejak R G and Mormann F 2013 Monitoring spike train synchrony. J. Neurophysiol. 109 1457–72

[44] Satuvuori E and Kreuz T 2018 Which spike train distance is most suitable for distinguishing rate and temporal coding? J. Neurosci. Methods 299 22–33

[45] Cubero R J, Marsili M and Roudi Y Finding informative neurons in the brain using Multi-Scale Relevance

[46] Azarfar A, Calcini N, Huang C, Zeldenrust F and Celikel T 2018 Neural coding: A single neuron’s perspective. Neurosci. Biobehav. Rev. 94 238–47

[47] Miceli S, Nadif Kasri N, Joosten J, Huang C, Kepser L, Proville R, Selten M M, van Eijs F, Azarfar A, Homberg J R, Celikel T and Schubert D 2017 Reduced Inhibition within Layer IV of Sert Knockout Rat Barrel Cortex is Associated with Faster Sensory Integration. Cereb. Cortex 27 933–49

[48] Huang C, Resnik A, Celikel T and Englitz B 2016 Adaptive spike threshold enables robust and temporally precise neuronal encoding. PLoS Comput. Biol. 12 e1004984

[49] Avitan L and Goodhill G J 2018 Code under construction: neural coding over development. Trends Neurosci. 41 599–609

[50] Kole K, Scheenen W, Tiesinga P and Celikel T 2018 Cellular diversity of the somatosensory cortical map plasticity. Neurosci. Biobehav. Rev. 84 100–15

[51] Quiroga R Q 2012 Spike sorting. Curr. Biol. 22 R45–6

[52] Harris K D, Quiroga R Q, Freeman J and Smith S L 2016 Improving data quality in neuronal population recordings. Nat. Neurosci. 19 1165–74

[53] Ekanadham C, Tranchina D and Simoncelli E P 2014 A unified framework and method for automatic neural spike identification. J. Neurosci. Methods 222 47–55

[54] Tiganj Z and Mboup M 2011 A non-parametric method for automatic neural spike clustering based on the non-uniform distribution of the data. J. Neural Eng. 8 066014

[55] Einevoll G T, Franke F, Hagen E, Pouzat C and Harris K D 2012 Towards reliable spike-train recordings from thousands of neurons with multielectrodes. Curr. Opin. Neurobiol. 22 11–7

[56] Chung J E, Magland J F, Barnett A H, Tolosa V M, Tooker A C, Lee K Y, Shah K G, Felix S H, Frank L M and Greengard L F 2017 A fully automated approach to spike sorting. Neuron 95 1381–1394.e6

[57] Mokri Y, Salazar R F, Goodell B, Baker J, Gray C M and Yen S-C 2017 Sorting Overlapping Spike Waveforms from Electrode and Tetrode Recordings. *Front*. Neuroinformatics 11 53

[58] Mahmud M and Vassanelli S 2016 Processing and Analysis of Multichannel Extracellular Neuronal Signals: State-of-the-Art and Challenges. Front. Neurosci. 10 248

[59] Franke F, Pröpper R, Alle H, Meier P, Geiger J R P, Obermayer K and Munk M H J 2015 Spike sorting of synchronous spikes from local neuron ensembles. J. Neurophysiol. 114 2535–49

[60] Hilgen G, Sorbaro M, Pirmoradian S, Muthmann J-O, Kepiro I E, Ullo S, Ramirez C J, Puente Encinas A, Maccione A, Berdondini L, Murino V, Sona D, Cella Zanacchi F, Sernagor E and Hennig M H 2017 Unsupervised Spike Sorting for Large-Scale, High-Density Multielectrode Arrays. Cell Rep. 18 2521–32

[61] Rossant C, Kadir S N, Goodman D F M, Schulman J, Hunter M L D, Saleem A B, Grosmark A, Belluscio M, Denfield G H, Ecker A S, Tolias A S, Solomon S, Buzsaki G, Carandini M and Harris K D 2016 Spike sorting for large, dense electrode arrays. Nat. Neurosci. 19 634–41

[62] Lee J, Carlson D, Shokri H, Yao W, Goetz G, Hagen E, Batty E, Chichilnisky E J, Einevoll G and Paninski L 2017 YASS: yet another spike sorter BioRxiv

[63] Carlson D E, Vogelstein J T, Qisong Wu, Wenzhao Lian, Mingyuan Zhou, Stoetzner C R, Kipke D, Weber D, Dunson D B and Carin L 2014 Multichannel electrophysiological spike sorting via joint dictionary learning and mixture modeling. IEEE Trans Biomed Eng 61 41–54

[64] Magland J F and Barnett A H 2015 Unimodal clustering using isotonic regression: ISO-SPLIT

[65] Calabrese A and Paninski L 2011 Kalman filter mixture model for spike sorting of non-stationary data. J. Neurosci. Methods 196 159–69

[66] Redish A D 2017 MClust

[67] Carlson D E, Rao V, Vogelstein J T and Carin L 2013 Real-Time Inference for a Gamma Process Model of Neural Spiking Advances in Neural Information Processing Systems 26 (NIPS)

[68] Rutishauser U, Schuman E M and Mamelak A N 2006 Online detection and sorting of extracellularly recorded action potentials in human medial temporal lobe recordings, in vivo. J. Neurosci. Methods 154 204–24

[69] Quiroga R Q, Nadasdy Z and Ben-Shaul Y 2004 Unsupervised spike detection and sorting with wavelets and superparamagnetic clustering. Neural Comput. 16 1661–87

[70] Harris K D, Henze D A, Csicsvari J, Hirase H and Buzsáki G 2000 Accuracy of tetrode spike separation as determined by simultaneous intracellular and extracellular measurements. J. Neurophysiol. 84 401–14

[71] Lefebvre B, Yger P and Marre O 2016 Recent progress in multi-electrode spike sorting methods. J. Physiol. Paris 110 327–35

[72] Schmitzer-Torbert N and Redish A D 2004 Neuronal activity in the rodent dorsal striatum in sequential navigation: separation of spatial and reward responses on the multiple T task. J. Neurophysiol. 91 2259–72

[73] Schmitzer-Torbert N, Jackson J, Henze D, Harris K and Redish A D 2005 Quantitative measures of cluster quality for use in extracellular recordings. Neuroscience 131 1–11

[74] Shan K Q, Lubenov E V and Siapas A G 2017 Model-based spike sorting with a mixture of drifting t-distributions. J. Neurosci. Methods 288 82–98

[75] Liu X, Wan H and Shi L 2014 Quality metrics of spike sorting using neighborhood components analysis. Open Biomed. Eng. J. 8 60–7

[76] Pillow J W, Shlens J, Chichilnisky E J and Simoncelli E P 2013 A model-based spike sorting algorithm for removing correlation artifacts in multi-neuron recordings. PLoS ONE 8 e62123

[77] Franke F, Natora M, Boucsein C, Munk M H J and Obermayer K 2010 An online spike detection and spike classification algorithm capable of instantaneous resolution of overlapping spikes. J. Comput. Neurosci. 29 127–48

[78] Celikel T and Sakmann B 2007 Sensory integration across space and in time for decision making in the somatosensory system of rodents. Proc Natl Acad Sci USA 104 1395–400

[79] Azarfar A, Zhang Y, Alishbayli A, Miceli S, Kepser L, van der Wielen D, van de Moosdijk M, Homberg J, Schubert D, Proville R and Celikel T 2018 An open-source high-speed infrared videography database to study the principles of active sensing in freely navigating rodents. Gigascience 7

[80] Heckman J, McGuinness B, Celikel T and Englitz B 2016 Determinants of the mouse ultrasonic vocal structure and repertoire. Neurosci. Biobehav. Rev. 65 313–25

[81] Górska U, Rupp A, Boubenec Y, Celikel T and Englitz B 2018 Evidence Integration in Natural Acoustic Textures during Active and Passive Listening. Eneuro 5

[82] Heckman J J, Proville R, Heckman G J, Azarfar A, Celikel T and Englitz B 2017 High-precision spatial localization of mouse vocalizations during social interaction. Sci. Rep. 7 3017

[83] Zonooz B, Arani E, Körding K P, Aalbers P A T R, Celikel T and Van Opstal A J 2019 Spectral weighting underlies perceived sound elevation. Sci. Rep. 9 1642

[84] Freudenberg F, Resnik E, Kolleker A, Celikel T, Sprengel R and Seeburg P H 2016 Hippocampal GluA1 expression in Gria1-/- mice only partially restores spatial memory performance deficits. Neurobiol. Learn. Mem. 135 83–90

[85] Freudenberg F, Marx V, Seeburg P H, Sprengel R and Celikel T 2013 Circuit mechanisms of GluA1-dependent spatial working memory. Hippocampus 23 1359–66

[86] Seib D R M, Corsini N S, Ellwanger K, Plaas C, Mateos A, Pitzer C, Niehrs C, Celikel T and Martin-Villalba A 2013 Loss of Dickkopf-1 restores neurogenesis in old age and counteracts cognitive decline. Cell Stem Cell 12 204–14

[87] da Silva Lantyer A, Calcini N, Bijlsma A, Kole K, Emmelkamp M, Peeters M, Scheenen W J J, Zeldenrust F and Celikel T 2018 A databank for intracellular electrophysiological mapping of the adult somatosensory cortex. Gigascience 7

[88] Lim J and Celikel T 2019 Real-time contextual feedback for close-loop control of navigation. J. Neural Eng.

[89] Voigts J, Sakmann B and Celikel T 2008 Unsupervised whisker tracking in unrestrained behaving animals. J. Neurophysiol. 100 504–15

[90] Voigts J, Herman D H and Celikel T 2015 Tactile object localization by anticipatory whisker motion. J. Neurophysiol. 113 620–32

[91] Frehlick Z, Williams I and Constandinou T G 2016 Improving neural spike sorting performance using template enhancement 2016 *IEEE Biomedical Circuits and Systems Conference (BioCAS)* 2016 IEEE Biomedical Circuits and Systems Conference (BioCAS) (IEEE) pp 524–7

